# Single-cell RNA-seq based elucidation of the antidepressant hyperforin biosynthesis *de novo* in St. John’s wort

**DOI:** 10.1101/2024.01.24.577018

**Authors:** Song Wu, Ana Luisa Malaco Morotti, Jun Yang, Ertao Wang, Evangelos C. Tatsis

## Abstract

Hyperforin is the compound responsible for the effectiveness of St. John’s wort (*Hypericum perforatum*) as an antidepressant, but its biosynthesis remains unknown. Gene discovery based on co-expression analysis of bulk RNA-sequencing data or genome mining failed to discover the missing steps in hyperforin biosynthesis. Here we sequenced the tetraploid *H. perforatum* genome. By single-cell RNA-seq, we identified a distinct type of cells, Hyper cells, wherein hyperforin biosynthesis *de novo* takes place. Through pathway reconstitution in yeast and tobacco, we identify and characterize four transmembrane prenyltransferases to resolve hyperforin biosynthesis. The hyperforin polycyclic scaffold is created by a reaction cascade involving an irregular isoprenoid coupling and a tandem cyclization. Our findings reveal how and where hyperforin is biosynthesized that enables synthetic-biology reconstitution of the complete pathway. These results deepen our comprehension of specialized metabolism at the cellular level, and we anticipate acceleration of pathway elucidation in plant metabolism.

## Main text

St. John’s wort (SJW, *Hypericum perforatum*, Hypericaceae) is a high valued medicinal plant with annual global sales exceeding US$6 billion^1^. In classical antiquity, extracts from aerial SJW organs were used to treat skin ailments due to antiseptic and anti-inflammatory properties^2^, while nowadays are prescribed to treat mild to moderate depression, associated with fewer adverse reactions over common antidepressants^3,4^. Its primary potent bioactive, hyperforin, acts similarly to traditional antidepressants by inhibiting neurotransmitter re-uptake like serotonin, though via a different mechanism^5–9^. Unlike conventional antidepressants, hyperforin induces Ca^2+^ and Na^+^ by uniquely activating TRPC6 receptors, disrupting cation gradients and reducing neurotransmitter uptake. This unique process makes hyperforin a promising agent for treating depression^10^.

*H. perforatum* is endemic in Eurasia and North Africa and introduced in the Americas and Oceania by human migration. It originates from an ancient hybridization event in Siberia between *H. maculatum* and *H. attenuatum*^11^. Typically tetraploid, with a chromosome number of 2n = 4x = 32^11^, the plant also reproduces asexually through facultative apomixis, leading to populations with diploid, triploid and hexaploid cytotypes^12^. Besides hyperforin, SJW also biosynthesizes diverse specialized metabolites including essential oils rich in complex terpenoids^13^, flavonoids, biflavonoids, phenolic acids and naphtodianthrones^14^.

Hyperforin is the first polycyclic polyprenylated acylphloroglucinol (PPAP) to be isolated^15^ (Figure 1A), a diverse class of natural products with various biological activities that are taxonomic markers for the family Hypericaceae^16–18^. Hyperforin’s unique structure features an isoprenyl group at C8 (Figure 1A), thus poses a challenging task for enantioselective synthesis, inspiring chemists to pursue novel organic-synthesis strategies^16–18^. Such synthesis routes involve multiple steps can result in total yield of 1.64%^19^. Despite hyperforin’s significance, its biosynthesis *de novo* is unresolved. With reconstitution of the pathway in microbial or plant synthetic-biology systems, obtaining hyperforin from such sources is a sustainable and environmentally superior strategy to isolate this valuable pharmacological agent.

**Figure 1.**
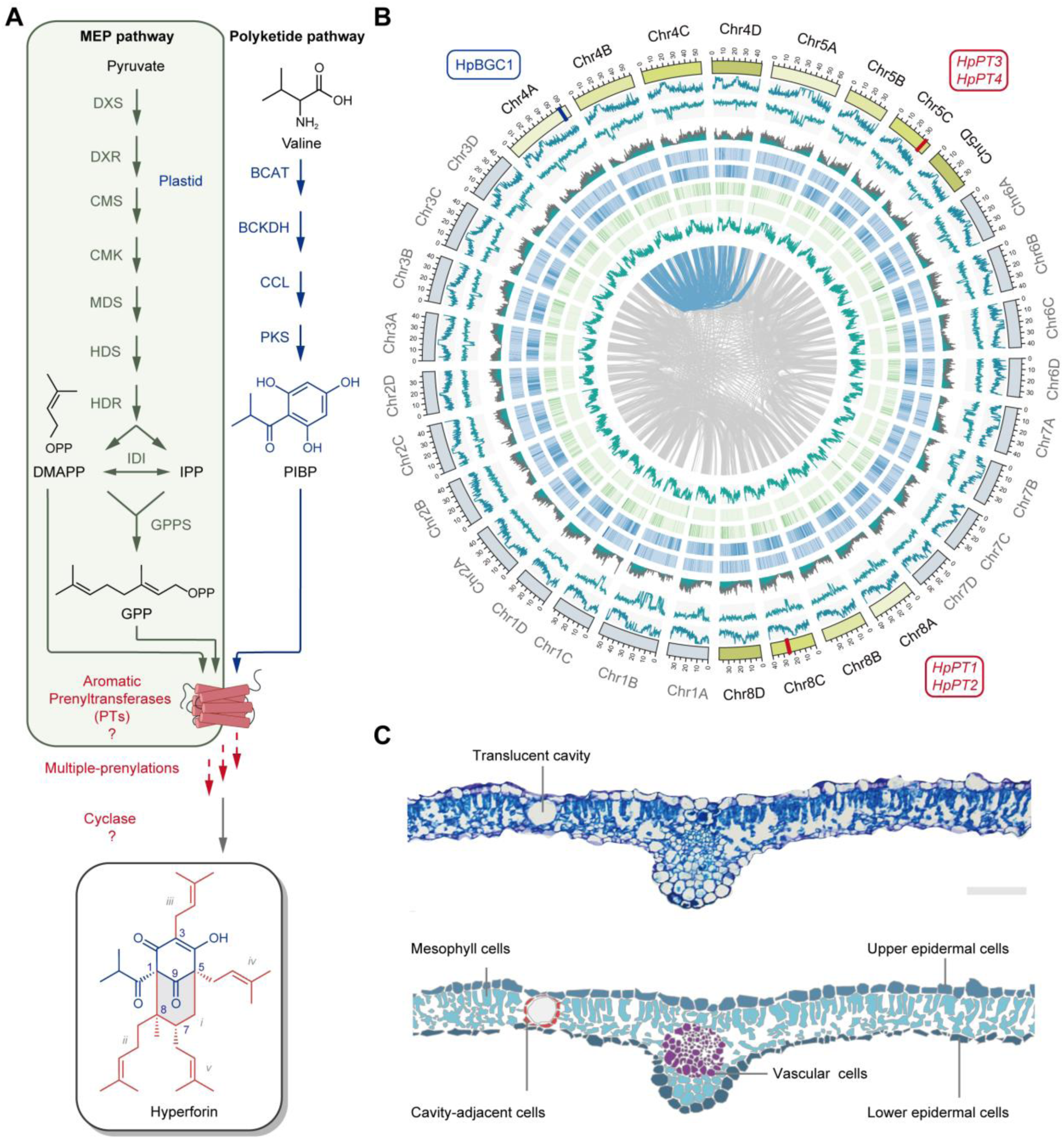
Hyperforin biosynthesis, translucent cavities and genomic architecture of *Hypericum perforatum*. **A.** Overview of the proposed model of hyperforin biosynthesis *de novo* in St. John’s wort. The pathway involves three metabolic modules. The first module involves the biosynthesis of polyketide phloroisobutyrophenone (PIBP) from valine^20^. The plastid-localized Methylerythritol 4-phosphate (MEP) pathway biosynthesizes prenyl units like dimethylallyl-diphosphate (DMAPP) and geranyl-diphosphate (GPP)^21^. The last module is hypothesized to involve prenylation of PIBP by a plastid-localized transmembrane aromatic prenyltransferase(s) and an enigmatic cyclization involving the formation of a C1–C8 bond^21–23^. **B.** Cross-section of *H. perforatum* leaf, highlighting the translucent pockets. Top: A tissue section stained with tolonium chloride, with a schematic depiction (below) depicting the spatial profile of the indicated cell types. Scale bar = 100 μm. **C.** Overview of *H. perforatum* genome assembly. The CIRCOS plot represents the features of the *H. perforatum* genome (size 1.39 Gb). From inner to outer layers: the gray links represent intraspecies syntenic relationships, densities of simple sequence repeats, rRNA, tRNA, snRNA, miRNA, genes, GC content, and transposable elements (TE) in 100-kb sliding windows, respectively. In the outer layer, bars represent individual chromosomes. Blocks marked in blue on Chr4A (*HpBGC1*), in red on Chr5C (*HpPT3– 4*), and Chr8C (*HpPT1–2*) indicate metabolic gene locations involved in hyperforin biosynthesis. Blue links highlight the syntenic relationships of *HpBGC1* copies among different chromosomes.

To date, details of the biosynthesis of specialized metabolites in SJW, and for the majority of the hyperforin biosynthesis steps, are sparse. We have characterized three enzymes, namely a branched-chain amino-acid dehydrogenase (BCKDHA), a CoA ligase (CCL) and an aromatic polyketide synthase (PKS) from two biosynthetic gene clusters (*HpBGC1–2*) that encode the first three steps dedicated to biosynthesis of the polyketide hyperforin precursor — phloroisobutyrophenone (PIBP)^20^ (Figure 1A). Provision of ^13^C-labeled substrates to *H. perforatum* seedlings revealed that the rest of the hyperforin carbon atoms comprise five isoprenoid units derived from the plastid-localized methyl-*D*-erythritol-4-phosphate (MEP) pathway^21^ (Figure 1A). Chemical logic suggests that the isoprenoid units derived from the plastid localized MEP pathway^21–23^ are adjoined to the polyketide PIBP by electrophilic substitution, involving an additional ring closure to biosynthesize the bicyclo[3.3.1]nonane scaffold (Figure S1) of hyperforin. Transfer of isoprenoid moieties from pyrophosphate donor substrates to aromatic acceptor substrates in plants is typically catalyzed by membrane-bound aromatic prenyltransferases ^24–27^.

High-quality genome assemblies are of paramount importance for complete pathway elucidation of metabolites with complicated structures^28,29^. Two fragmented and incomplete annotations of a diploid *H. perforatum* genome are available ^30,31^, but these are insufficient for hyperforin pathway elucidation (Figure S2).

Here, to fully elucidate the hyperforin biosynthetic pathway, we sequenced the tetraploid genome of *H. perforatum* and generated single-cell atlases for leaves and flowers. Based on mapping of previously characterized upstream genes, we identified a new type of cells — Hyper cells — wherein biosynthesis *de novo* of hyperforin occurs. By using two synthetic-biology platforms for heterologous expression (yeast and tobacco), we discover and characterize four new UbiA-family prenyltransferases that complete the hyperforin biosynthesis pathway. The last step for hyperforin biosynthesis involves an unusual prenyltransferase catalyzing an irregular branching prenylation and a rare tandem cyclization.

## Results

### Sequencing and assembly of the tetraploid *Hypericum perforatum* genome

To enable the discovery of the missing steps in hyperforin biosynthesis, we generated transcriptome and genome-sequence resources *de novo* for *H. perforatum* using a suite of Illumina, PacBio HiFi and Hi-C sequencing (Table S1). We used Illumina paired-end short-read sequencing to survey the genome, and based on k-mer analysis, the genome size was estimated as 1.71 Gb (Table S2). PacBio HiFi long-read sequencing was mainly used for genome assembly in conjunction with Illumina sequencing for contig error correction, resulting in a 1.54-Gb (Table S3) draft assembly comprising 3,710 scaffolds and an N50 value 42.44 Mb. A Hi-C (*in vivo* fixation of chromosomes) library refined the first version of the reference genome and the scaffolds aligned into 32 groups hereafter referred to as chromosomes (Figure 1B, Table S4, Figure S3), which represented a tetraploid accession with a haploid chromosome number of 8. Coverage of the final assembly was evaluated by Illumina short-read mapping, CEGMA and BUSCO (Tables S5–7) that indicate a high-quality assembly with 97.6% completeness. A total of 98,005 genes (Table S8) were annotated by a pipeline combining *de novo* and homology-based predictions, with bulk RNA-seq data^32^. Transposable elements account for 56.95% of the genome (Table S9).

Functional annotation of genes was performed by searching against publicly available databases including SwissProt, NCBI-NR, Pfam, KEGG and InterPro. The best match from each database was assigned to the corresponding protein-coding gene. Following successful anchoring of 32 chromosomes and phasing of four sub-genomes by Hi-C (Figure S4A), we mapped the location of upstream genes within the known biosynthetic gene cluster *HpBGC1*^20^. *HpBGC1* is located on Chr4A within a topologically associating domain (TAD)^33^ (Figure S4B), highlighting the co-regulation of the cluster’s genes for proper biosynthesis of the polyketide precursor PIBP. Intraspecies syntenic analysis revealed the presence of five allele counterparts of *HpBGC1* on Chr4A-D and on the non-homologous chromosome Chr5A (Figure S5A). These copies of *HpBGC1a–e* are not identical, but bear variation in metabolic-gene co-linearity, gene-copy numbers and tandem duplications (Figure S5B). By contrast, *HpBGC1d* in Chr4D has undergone substantial gene losses and retained only one predicted *PKS*. These *HpBGC1* polymorphisms are possibly the result of segmental re-arrangements within the tetraploid genome.

### Genome mining and co-expression analysis to identify aromatic prenyltransferases

Several instances are reported of three or more genes comprising the same biosynthetic pathway are physically co-localized, thus genome mining can aid gene discovery^33–36^. Because the polyketide-module biosynthesis genes clustered within *HpBGC1*^20^, clustering may well also be the case for the downstream genes^32,37,38^. Based on functional annotation, 405 genes were predicted to be UbiA-family aromatic prenyltransferases. By mining the *H. perforatum* genome, we identified a cluster comprising 11 annotated aromatic-prenyltransferase genes bearing high homology to known enzymes acting in biosynthesis of prenylated xanthones^39^. A blueprint of prenylated-xanthone metabolism in *Hypericum* species^40–42^ indicates analogies with the proposed model of hyperforin biosynthesis involving the recruitment of dedicated CoA ligases, type-III polyketide synthases and aromatic prenyltransferases that differ in the starting polyketide unit and the role of oxidative cyclizations in the xanthone pathway^43,44^ (Figure S6). Toward determining their functionality and potential contribution to hyperforin biosynthesis in the absence of available functional-characterization approaches in *H. perforatum*, the 11 predicted prenyltransferase genes were individually cloned and co-expressed in yeast together with the known CoA ligase (*HpCCL2*) and polyketide synthase (*HpPKS2*) from *HpBGC1*^20^. No prenylated products were detected by LC–MS analysis, indicating that these prenyltransferases do not participate in hyperforin biosynthesis.

Genes in plant specialized-metabolite biosynthesis are usually co-regulated as part of the same biological pathway, thus discovery of unknown metabolic genes in such pathways is usually based on co-expression analysis of bulk RNA-seq data across plant samples (e.g. various tissues or different treatments)^28,45–47^. Co-expression analysis with upstream genes from *HpBGC1* as bait did not yield likely prenyltransferase candidates with high correlation efficiency^45,47^ (Pearson Correlation Coefficient > 0.90) among the 405 annotated prenyltransferases (Figure S7).

### Single-cell RNA sequencing enables discovery of missing hyperforin biosynthetic genes

The eponym *perforatum* for St John’s wort is derived from the visible translucent spots that resemble perforations found on both leaves and flowers that also are present in certain other *Hypericum* species^48,49^ (Figure 1C). These translucent cavities are spherical cellular structures spanning the upper to lower epidermis delimited by two cell layers — an inner layer consisting of flattened secretory-like cells and an outer layer comprising turgid parenchymatous cells^50,51^. Hyperforin accumulates exclusively within these structures in *H. perforatum* leaves^51^. In contrast, hyperforin is not detectable in non-secretory tissues, indicating that the hyperforin present in the secretory structures originates exclusively from either the translucent cavities or from the immediately adjacent cells^51^. It remains unclear whether the complete hyperforin biosynthesis, or step/s in it, is active around the translucent cavities, or whether they simply serve as storage vessels.

High-throughput single-cell RNA sequencing (scRNA-seq) can distinguish cell types or cell populations based on cell-to-cell heterogeneity as regards gene expression^52–55^, but its mainstream exploitation in plant science is hindered by substantial technical barriers such as the preparation of protoplasts and the heterogeneity of plant tissues^52,54,55^. Despite this, scRNA-seq has been applied to cellular differentiation and communication^52,54^, while use in plant specialized metabolism is sparse^53,56^.

We hypothesize that since hyperforin biosynthesis correlates spatially with translucent cavities and their associated cell types (Figure 1C), then the expression signal of hyperforin biosynthesis genes might be concealed by noise typical of bulk RNA-seq, thus necessitating scRNA-seq to carefully examine the cellular architecture of *H. perforatum* leaves. To generate a single-cell transcriptome atlas of *H. perforatum* leaves, we successfully isolated viable protoplasts from young leaves to capture the cellular heterogeneity within the tissue. Droplet-based 10x Genomics was used for RNA-seq^57^ to capture gene-expression data at the single-cell level. Reads were mapped to the reference genome, quantifying gene-expression levels in individual cells with 92.2% of the reads mapped to the reference genome, thus validating the quality and accuracy of the transcripome sequence. We identified a total of 15,074 cells expressing a median of 1,405 genes across different cell types. To interpret this dataset and categorize the cell types, low-quality and doublet cells were removed, followed by normalization of the expression data to eliminate technical noise and standardization for downstream analysis. Principal component analysis (PCA) was used to reduce the dimensionality of scRNA-seq data and distill the complex gene-expression profiles, capturing the most significant variation among the assayed cells. To categorize the cells into clusters based upon shared transcriptome profiles, we applied the unbiased clustering algorithm uniform manifold approximation and projection (UMAP)^58^ to visualize and identify distinct cell populations (Figures 2A and S8).

**Figure 2.**
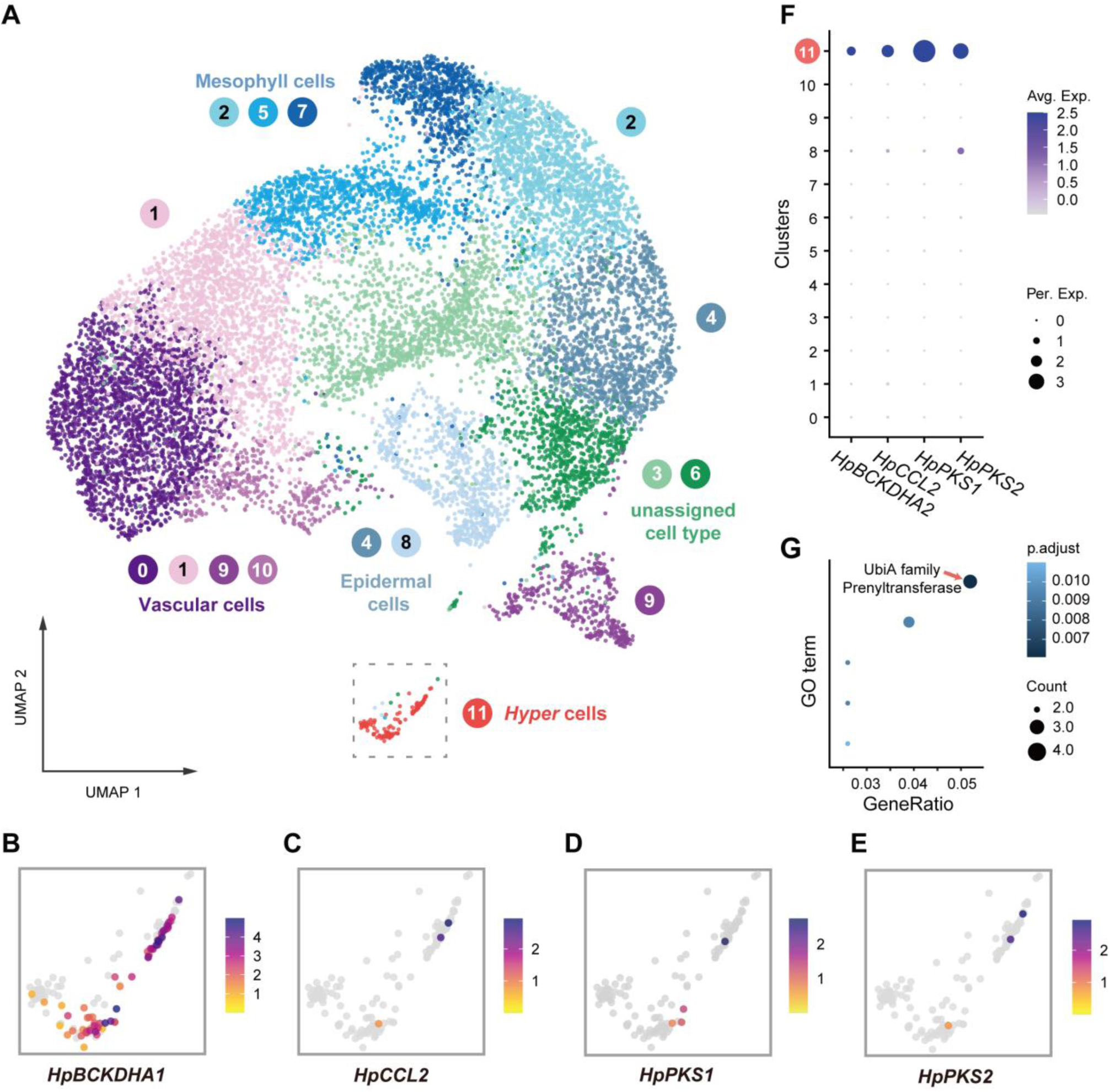
A single-cell atlas of a *H. perforatum* leaf and identification of Hyper cells. **A.** Dimensionally reduced UMAP visualization of single leaf cells with each color representing a distinct cluster. Twelve different cell clusters were identified. The four clusters of vascular cells (0, 1, 9 and 10) are visualized in shades of purple–pink; the three clusters of mesophyll cells (2, 5 and 7) are visualized in shades of blue–cyan; the two clusters of epidermal cells (4 and 8) in shades of blue–grey; and two clusters of unassigned cells (3 and 6) in shades of green. Cells of cluster 11, visualized in red and enclosed within a dashed box, are assigned as Hyper cells. Each dot represents one cell **B–E**. Close-up view of the expression profiles in Hyper cells of genes previously identified in hyperforin biosynthesis *HpBCKDHA1*, *HpCCL2*, *HpPKS1* and *HpPKS2* from *HpBGC1*. The heatmap color scale depicts the relative expression of each gene in each cell. **F**. Dot plot illustrating the predominant expression of metabolic genes in the polyketide module of hyperforin biosynthesis (*HpBCKDHA2*, *HpCCL2*, *HpPKS1* and *HpPKS2* from *HpBGC1*) across all clusters in this atlas. The color scale indicates the average scaled expression of each gene in each cell type. Dot sizes depict the fraction of cells wherein a given gene is expressed in a given cell type. **G**. GO enrichment analysis of cluster 11, revealing ‘UbiA-family prenyltransferase’ as the top-enriched protein family.

Annotation of cell types in a plant single-cell atlas is based on single or a set of marker genes. However, such markers are absent for *H. perforatum*. There is, however, an extensive database of genes from mainstream model plants used as cell-type markers^59–61^. Identification of functional orthologs between *A. thaliana* and *H. perforatum* by sequence alignment and homology are prone to false positives. Instead, synteny is one of the most reliable criteria for establishing the functional relationships between orthologous genes or genomic regions across different species^62^. To annotate the cell clusters with an unbiased method, we performed a fully automated annotation pipeline comprising: (1) compiling an extensive marker-gene dataset for *A. thaliana*^59^; (2) through syntenic analysis between *A. thaliana* and *H. perforatum*, we correlated the orthologous marker genes between the two species and generated a marker-gene dataset for *H. perforatum*; and (3) using the ScType^63^ platform, an automated cell-type annotation was performed based on a pool of markers (Figure S9). Twelve different cell types (Figures 2A and S8, Table S10) were identified with the cells from clusters 0, 1, 9, 10 annotated as vascular cells, the cells from clusters 2, 5, 7 are mesophyll cells, and cells from clusters 4 and 8 are epidermal cells. There were no adequate markers to support annotation for cells from clusters 3 and 6, while the minority cell-type cluster 11 was annotated with low confidence (i.e. low ScType score) as myrosin cells^59,64^.

Visualization and quantification of expression of known early hyperforin biosynthetic-pathway genes^20^ revealed that *HpBCKDHA1*, *HpBCKDHA2*, *HpCCL2*, *HpPKS1* and *HpPKS2* are distinctively and highly expressed in cluster 11 (Figure 2B–F). This indicates that biosynthesis of the precursor PIBP takes place predominantly in cell-cluster 11, and in parallel, raises the question whether genes encoding downstream steps are co-expressed in these cells. Toward addressing this, gene-ontology (GO) enrichment analysis^65^ found genes belonging to the UbiA prenyltransferase family are over-represented in this cluster (Figures 2G and S10). The isoprenoid carbon atoms in hyperforin are derived from the MEP plastidial pathway^21^ and UbiA-type transmembrane aromatic prenyltransferases are expected to catalyze the prenylation reactions by transferring isoprenoid units to PIBP. MEP pathway genes *HPDXS*, *HpHDS1*, *HpHDS2*, *HpHDR* and *HpGPPS* are all highly expressed in vascular cells (cluster 0) and in cluster 11 (Figure 3A,B). We therefore focused on aromatic prenyltransferases expressed in cell-cluster 11. Based on this profile, four genes annotated as aromatic prenyltransferases (*Hper_g146976*, *Hper_g147016*, *Hper_g96819* and *Hper_g96821*) were selected as candidates based on their high expression in cluster 11 (Figure 3B,C), and successfully cloned. All the selected candidate prenyltransferases have plastid transit peptides at the N-termini, which were removed for expression in yeast based on ChloroP predictions^66^.

**Figure 3.**
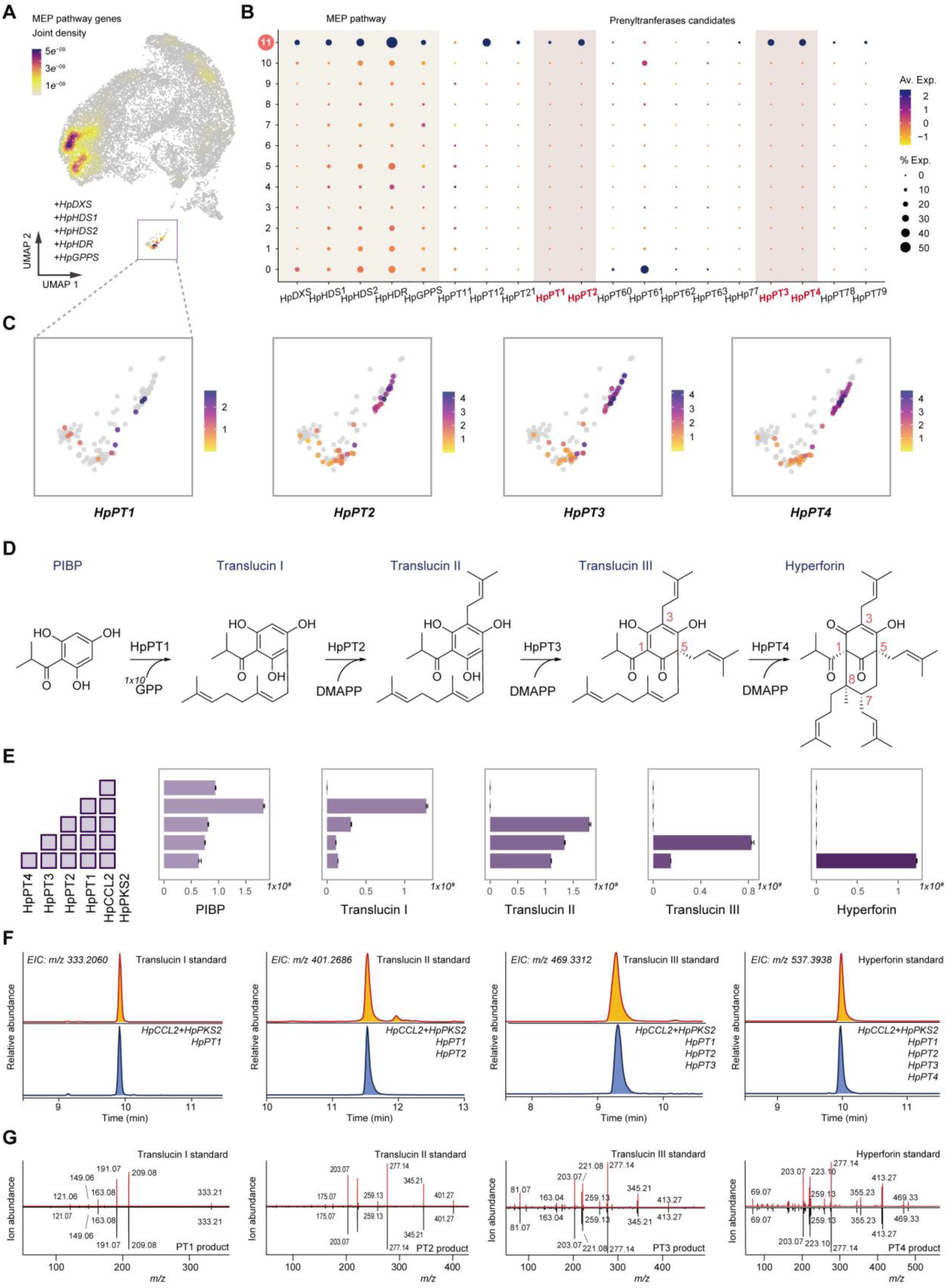
Elucidation of the catalytic steps to complete the formation of hyperforin. **A.** Overlaid gene expression of MEP pathway genes (*HpDXS*, *HpHDS1*, *HpHDS2*, *HpHDR* and *HpGPPS*) across the *H. perforatum* leaf single-cell atlas demonstrates high expression in cells clusters 0 (vascular) and 11 (Hyper cells). The heatmap color scale indicates the sum of expression from the 5 MEP genes in each cell. **B.** Dot plot highlighting elevated expression levels of MEP pathway genes in cluster 11, along with potential prenyltransferase (*PT*) candidate genes. Genes with confirmed activity are marked in red. The color scale indicates the average scaled expression of each gene in each cell type. Dot sizes reflect the fraction of cells wherein a given gene is expressed in a given cell type. **C.** Cell-level expression profiles of prenyltransferases (*HpPT1*, *HpPT2*, *HpPT3* and *HpPT4*) in Hyper cells (cluster 11). The heatmap color scale shows the expression of each gene in each cell. **D.** The sequential prenylation and tandem cyclization steps leading to hyperforin formation from phloroisobutyrophenone (PIBP), based on the verified activity of prenyltransferases HpPT1, HpPT2, HpPT3 and HpPT4. The intermediates translucins I–III were chemically synthesized and detected in *H. perforatum* extracts (Figures S11 and S13–15). **E.** LC–MS peak area of the exact ion mass [M + H]^+^ for each intermediate produced in yeast after co-expression of the indicated genes. Boxes to the left of the plots denote combinations of biosynthetic genes in each co-expression experiment. Data were obtained from three independent biological replicates and the bars represent the mean ± SE. **F.** Comparison of extracted ion chromatogram (EIC) of each PIBP prenylated product from *in vivo* co-expression of biosynthetic genes (in red–gold) with EICs from corresponding synthetic standards (translucins I–III) and hyperforin (in black–blue). **G.** Comparative MS/MS fragmentation patterns (MS^2^ spectra) of prenylated products and standards from panel F.

To screen the candidate prenyltransferases involved in hyperforin biosynthesis (Figure 3D, E), we assembled the PIBP precursor module in *Saccharomyces cerevisiae*. The CoA ligase HpCCL2 and the type-III PKS HpPKS2^20^ were cloned into pESC-Leu and transcripts lacking the corresponding sequence encoding plastidial transit peptides of the four candidate prenyltransferases were cloned into pESC-His and transformed into the isoprenoid-rich yeast strain DD104^67^. After galactose induction, the culture was extracted with ethyl acetate and analyzed by LC–MS. Only the prenyltransferase encoded by *Hper_g146976* demonstrated activity towards PIBP with a new peak eluting at 9.9 min (Figures 3F and S11) under the extracted ion chromatogram (EIC) mode at *m*/z 333.2060, suggesting the enzyme uses a geranyl-diphosphate molecule as prenyl donor and attaches the geranyl chain to PIBP. The geranyl-PIBP synthesized standard elutes at 9.9 min with *m/z* = 330.2060 in positive mode, which aligned with the peak of the Hper_g146976 product (Figure 3F) and is in agreement with the MS^2^ spectrum (Figures 3G and S11). We conclude that the prenyltransferase encoded by *Hper_g146976* acts on PIBP and catalyzes the first prenylation step in hyperforin biosynthesis by way of a geranyl transferase. It is therefore assigned HpPT1 and its product as translucin I. Further analysis of enzyme-assay LC–MS data showed that HpPT1 uses only GPP as a prenyl donor and not DMAPP (Figure S12).

To elucidate the outstanding biosynthesis steps involving PIBP prenylations, we cloned the three remaining candidate prenyltransferase genes into the second cloning site of pESC-His and assayed their activity in yeast. EtOAc extracts from galactose-induced strains were analyzed by LC–MS and a new peak at EIC *m/z* 401.2686 was detected at 11.6 min (Figures 3E,F and S13) in a strain expressing *Hper_g147016.* By comparison with the translucin-II synthetic standard and alignment of chromatograph (based on retention time, Figure 3F) and MS data (MS and MS^2^, Figure 3G), we concluded that *Hper_g147016* encoding HpPT2 catalyzes the second prenylation in hyperforin biosynthesis.

To discover further the remaining prenylation steps, we cloned the remaining two candidates into pESC-Ura and assayed their activity using the same workflow. When we co-expressed *Hper_g96819* with HpCCL2, HpPKS2, HpPT1 and HpPT2, a new peak appeared in the LC–MS chromatogram of EIC *m/z* of 469.3312 at 9.4 min (Figures 3D–F and S14). This new product was identified as translucin III based on comparison of retention time, MS and MS^2^ spectrum with a synthetic standard (Figures 3D–F and S14), indicating the geminal prenylation on C5 (Figure 3D) as HpPT1 and the de-aromatization of the polyketide ring. Therefore, Hper_g96819 was assigned HpPT3 that catalyzes the third prenylation step on the scaffold of PIBP polyketide.

Based on chemical intuition (Figure 3D), a prenylation and a cyclization step are expected for the transformation of translucin III to hyperforin. To determine whether the only remaining candidate gene encodes such functionalities, *Hper_g96821* was cloned into the second site of pESC-Ura and co-expressed with HpCCL2, HpPKS2 and HpPT1–3. Hper_g96821 is enzymatically active, transforming translucin III to hyperforin, with the product eluting at 10.0 min (the same time as the hyperforin standard), and the MS and MS^2^ spectra are identical to those of hyperforin (Figures 3D– G and S15). Hper_g96821 catalyzes the fourth sequential prenylation step and was thus named HpPT4.

Although HpPT4 is a UbiA prenyltransferase, it does not apparently catalyze a typical Friedel– Crafts alkylation^24,26^ as an aromatic prenyltransferases are expected to, but instead acts as an irregular prenyltransferase catalyzing the 1′-2 coupling (branching)^25,68^ between two isoprenyl units (Figure S1A). Moreover, it appears that HpPT4 does not only catalyze the transfer of a prenyl unit to translucin III, but also catalyzes bond formation between C(1) and C(8), resulting in cyclization and biosynthesis of the iconic bicyclo[3.3.1]nonane (Fig. S1A) scaffold of PPAPs^17,18^. Such cyclizations generating complex polycyclic structures are hallmarks of natural-product biosynthesis (Figure S1B)^69^.

To ensure host enzyme activities do not bias the functional characterization of candidates^70^, we assayed the activity of full-length HpPT1–4 by transient expression *Nicotiana benthamiana* leaves by co-expression with *HpBCKDHA1*, *HpCCL2* and *HpPKS2* followed by LC–MS analysis of leaf methanol extracts. Hyperforin was detected *in planta* only when all the enzymes in the pathway were expressed (Figure 4A). This supports the functional characterization in yeast and safeguards the proposed role of HpPT4 as a hyperforin synthase to transfer a prenyl group and catalyze the cyclization step to complete hyperforin biosynthesis.

**Figure 4:**
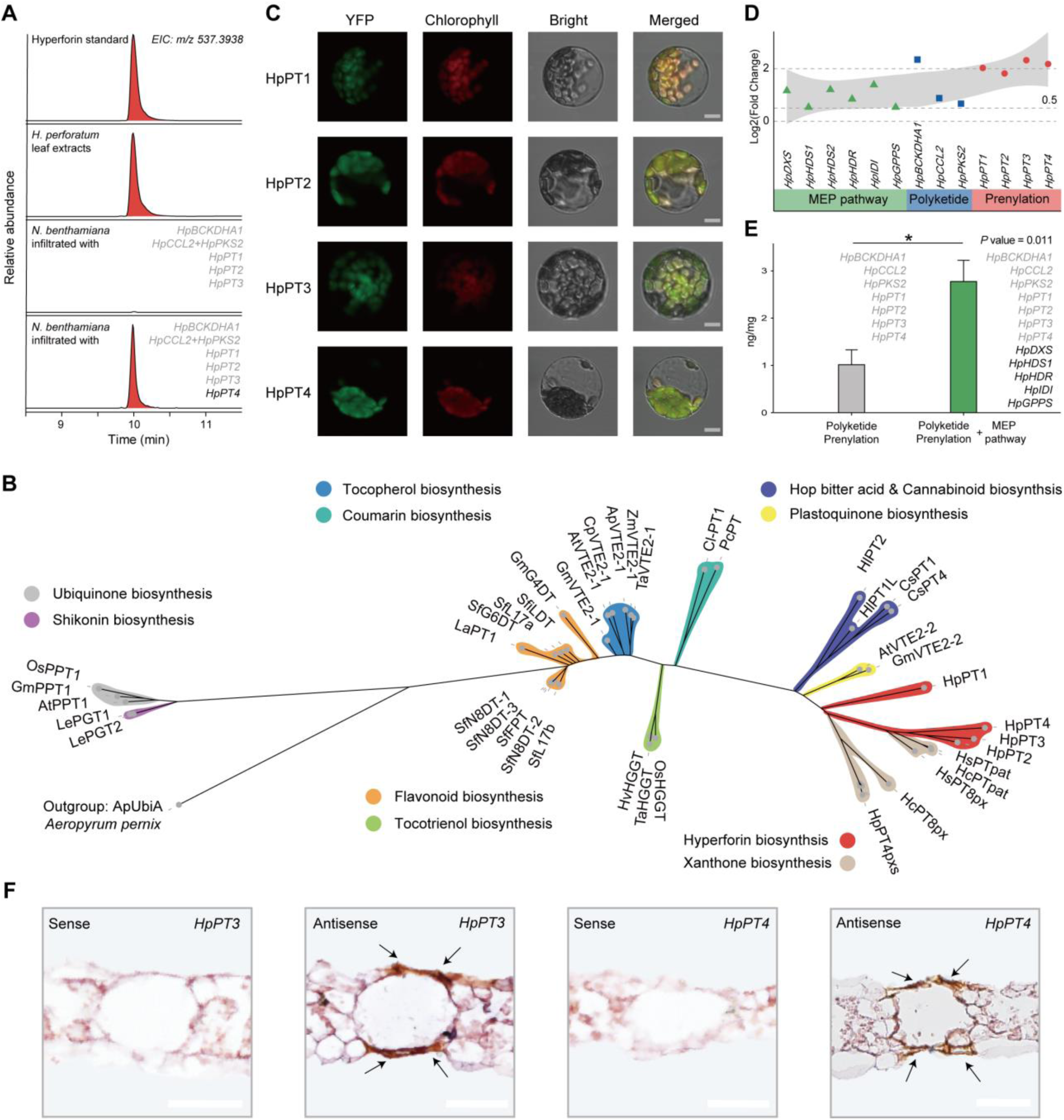
Plastids are prenyl-unit donors for hyperforin biosynthesis in *H. perforatum*. **A.** Analysis of transient co-expression of *HpBCKDHA1*, *HpCCL2*, *HpPKS2*, *HpPT1*, *HpPT2*, *HpPT3* and *HpPT4* in tobacco for biosynthesis of hyperforin by LC–MS. Extracted ion chromatograms (EIC m/z 537.3938) from LC-MS analysis of a hyperforin standard and *H. perforatum* extract are shown for comparison. **B.** Phylogenetic tree of characterized aromatic prenyl transferases from UbiA family from plants. The phylogeny was built from published, confirmed-active UbiA enzymes in plant metabolism and the prenyltransferases characterized in this study (highlighted in red shading) using 1,000 bootstrap values. UbiA from the fungus *Aeropyrum pernix* was used as the outgroup. The different color shades indicate the different pathways that the respective prenyltransferases are acting in. The phylogenetic analysis illustrating the close relation of prenyltransferases in hyperforin biosynthesis with the ones from plastid-localized plastoquinone biosynthesis (shaded in yellow). **C.** Subcellular localizations of HpPT1, HpPT2, HpPT3 and HpPT4 with predicted transit peptides fused at the C-terminus to YFP in transiently transgenic Arabidopsis leaf mesophyll protoplasts imaged by confocal laser-scanning microscopy. Scale bars = 10 µm **D.** Dot plot depicting the logarithmic differential enhanced expression of genes in leaves from the hyperforin biosynthetic modules MEP (green), polyketide/PIBP (blue) and prenylations (red) after 24 h of treatment with 400 µM methyl jasmonate. All the genes in hyperforin biosynthesis *de novo* show enhancement of their expression by methyl jasmonate. **E.** LC–MS quantification of hyperforin production by transient co-expression in tobacco (hyperforin (ng) per leaf sample dry weight (mg)) of the genes from polyketide and prenylation modules (in gray) and co-expression of the genes from MEP, polyketide and prenylation modules (in green). Data were obtained from five independent biological replicates. The bars represent the mean ± SE. **F.** RNA *in situ* hybridization reveals the localization of *HpPT3* and *HpPT4* expression in proximity to the translucent cavities in *H. perforatum* leaf cross-sections. Scale bars = 50 µm.

To further validate the proposed catalytic capacity of HpPT4 acting as branching 1′-2 prenyltransferase and cyclase, we tested the catalytic activity of HpPT4 against the translucin-III analog colupulone^67^ (Figure S16). We co-expressed HpPT4 with HpCCL2, HpPKS2, HlPT1L and HlPT2^67^ in yeast. LC–MS analysis of extracts showed that HpPT4 could transform colupulone to secohyperforin^71^ (8.0 min; *m/z* [M+H]^+^ 469.3312) in a similar manner as it transforms translucin III to hyperforin (Figure S16). To screen for additional prenyltransferases that have the same activity with HpPT1–4; we cloned, expressed and assayed HpPT12, HpPT21 and HpPT77 (Figures S11, S13– 15). No activity was observed.

### Hyperforin biosynthesis *de novo* involving all three modules takes place in Hyper cells

*HpPT1* and *HpPT2* are located close by on Chr8 (Figure S17A) and HpPT3 and HpPT4 are also located near one another on Chr5 (Figure S17B). Genomic proximity can imply that the two pairs are results of tandem duplications, however, the low sequence identity between these two pairs suggest this is not the case (Table S11). Phylogenetic analysis of annotated prenyltransferases based on amino-acid sequences reveals that HpPT1–4 are members of the HG clade of transmembrane plant aromatic prenyltransferases with plastid localization, similar to enzymes acting in the plastoquinone pathway (Figure 4B). HpPT1–4 fused at the C-terminus to YFP when transiently expressed in *Arabidopsis* protoplasts localize to plastids (Figure 4C), supporting conclusions from stable-isotope supplementation experiments indicating isoprenyl units are derived from the plastidial MEP pathway^21^.

Hyperforin biosynthesis can be divided in three metabolic modules involving the biosynthesis of the polyketide PIBP localized to the cytosol, the MEP pathway in plastids supplying prenyl donors (DMAPP, GPP), and plastid-localized transmembrane prenylation (Figure S18). We have previously shown that genes encoding PIBP biosynthetic enzymes are up-regulated 24 h after methyl jasmonate treatment^20^. Genes in the MEP pathway and *HpPT1–4* are also responsive to methyl jasmonate treatment, indicating co-regulation of the whole hyperforin biosynthetic pathway (Figures 4D and S19). To confirm the role of *H. perforatum* genes from the MEP pathway in supplying precursors for hyperforin biosynthesis, we transiently co-expressed the *H. perforatum* MEP genes (*HpDXS*, *HpDXR*, *HpHDS1*, *HpHDR*, *HpIDI* and *HpGPPS*) together with genes from the PIBP module and *HpPT1–4* in tobacco, resulting in over two-fold hyperforin production compared to co-expression of genes from polyketide and prenylation modules (Figure 4E).

Because all the newly identified genes involved in hyperforin biosynthesis are exclusively and highly expressed in cell-cluster 11, we therefore name them ‘Hyper cells’. Using Seurat^58^, we identified 15 high-confidence genes that can be used as markers for Hyper cells (Table S12 and Figure S20). Of these, *HpBCKDHA1*, *HpPT2*, *HpPT3* and *HpPT4* feature with expression at least 4-fold higher than other cell types. To visualize the position of Hyper cells in leaf cellular architecture, we performed RNA *in situ* hybridization using probes against *HpPT3* and *HpPT4*. *HpPT3* and *HpPT4* transcripts indicate Hyper cells are adjacent to translucent cavities (Figures 4F, S21 and S22).

### Biosynthesis *de novo* of hyperforin in flowers also takes place in Hyper cells

Because hyperforin strongly accumulates in both leaves and open flowers^20,72^, the discovery of Hyper cells and elucidation of hyperforin biosynthesis *de novo* in leaves raises the question about activity of the hyperforin biosynthetic pathway in flowers. To determine whether hyperforin biosynthesis *de novo* in floral organs is organized at the cellular level in the same way as leaves, protoplasts were isolated from flowers for scRNA-seq. The floral single-cell atlas reveals nine different cell clusters (Figure 5A and Table S13). Based on a customized annotation pipeline, we observed two types of epidermal cells (cell clusters 2 and 3), two types of vascular cells (cell clusters 1 and 4), one type of cells assigned as cortex cells (cell-cluster 7), one type of cells assigned as pollen cells (cell-cluster 8), and two type of unassigned cells (cell clusters 5 and 6). *HpPT1–4* are specifically and highly expressed in cell cluster 0 (Figure 5B*–*F) Moreover, expression of genes from all three modules of hyperforin biosynthesis (MEP, PIBP/polyketide and prenylation) predominantly occur in cell-cluster 0 in flowers (Figures 5B*–*F and S23).

**Figure 5:**
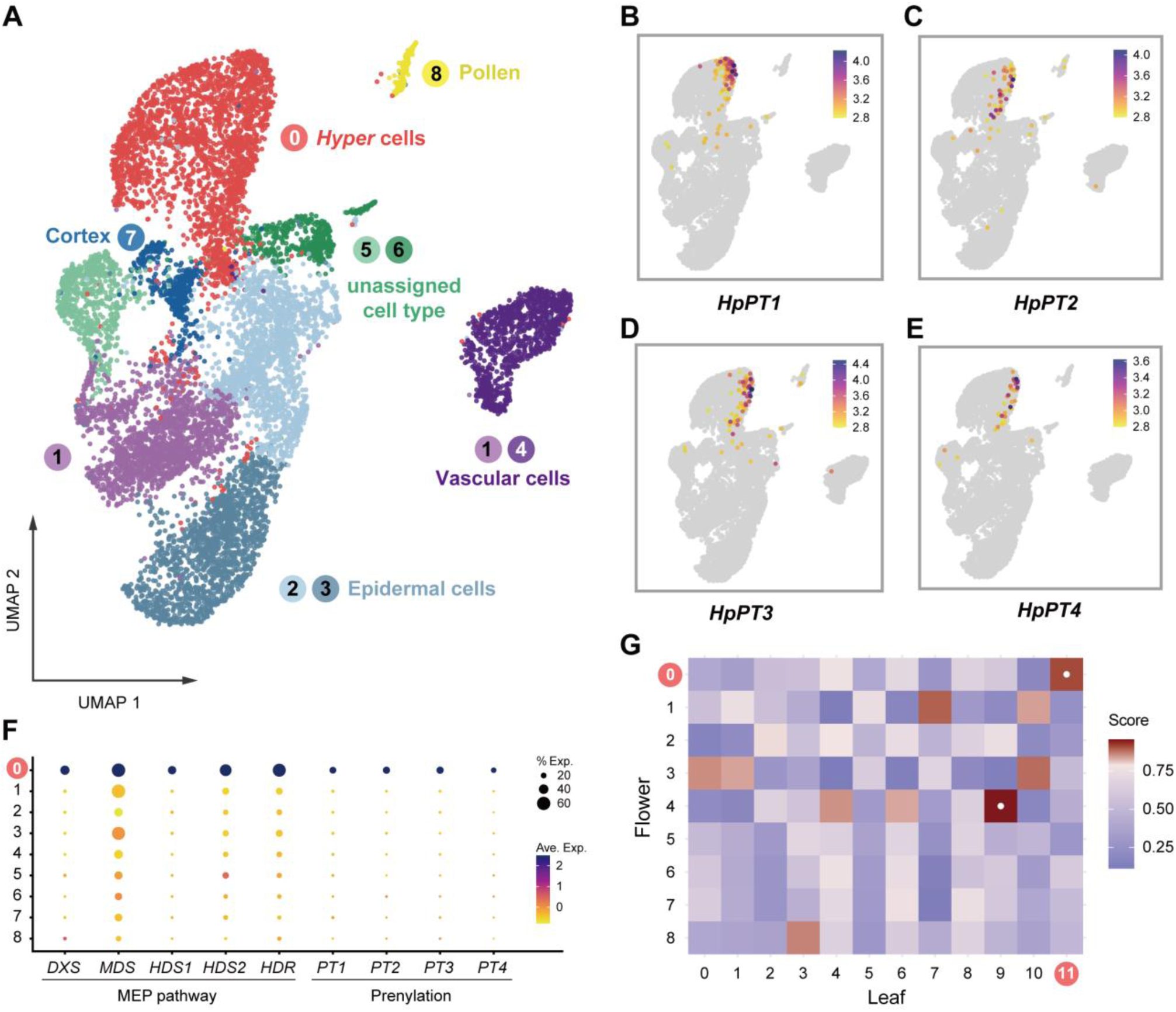
A single-cell atlas of *H. perforatum* flower reveals hyperforin biosynthesis in flowers. **A.** Dimensionally reduced UMAP visualization depicting various single-cell clusters within the flower, with each color denoting a distinct cluster **B–E**. Expression profiles of the indicated prenyltransferases (PTs) at the single-cell level within the flower. **F**. Dot plot showcasing the predominant expression patterns of metabolic genes involved in hyperforin biosynthesis *de novo* across identified clusters. **G**. Similarity matrix comparing cell clusters between the single-cell datasets of the flower and leaf, highlighting correlations with scores above 0.90 with white dots.

To explore the correlation and conservation of different cell types in leaves and flowers, especially regarding Hyper cells, we compared the respective scRNA-seq datasets using MetaNeighbor^73,74^, which through gene filtering, neighbor voting, and visualization, quantifies cell-type replicability, determines gene sets that contribute to cell-type identity and enables rapid identification of clusters with high similarity. A high level of similarity between cluster 11 from leaves with cluster 0 from flowers lead to confirmation of Hyper cells being present in the two tissues of St. John’s wort (Figure 5G).

## Discussion

By sequencing the genome of *H. perforatum* at chromosome level and generating single-cell atlases from leaves and flowers, we demonstrate how the combination of these approaches enabled the elucidation of hyperforin biosynthesis *de novo* and reconstitution of the complete pathway in yeast and tobacco. We anticipate these findings serve as a blueprint to elucidate the biosynthesis of other prominent bioactive PPAPs such as the anticancer molecule garcinol^75^ and neuroactive garsubellin A^76^ (Figure S1B). Our results illustrate how a UbiA-family aromatic prenyltransferase was recruited to catalyze the 1’-2 irregular prenylation reaction^25^ and also act as a terpene cyclase for the formation of the bicyclo[3.3.1]nonane scaffold found in hyperforin and certain other PPAPs in medicinal plants^17,18^. This complex bicyclic scaffold is a common structural feature among natural products with different biosynthetic origins (such as alkaloids, terpenoids, polyketides and meroterpenoids, Figure S1B)^46,77,78^ across plants and microorganisms. Hyperforin synthase (HpPT4) is one example of how nature’s enzymatic versatility permits different enzymes to biosynthesize this abundant yet complex bicyclic scaffold. Heterologous biosynthesis of hyperforin highlights potential synthetic-biology applications for production of this valuable molecule and other bioactive PPAPs.

Single-cell RNA sequencing serves as a powerful, alternative approach for gene discovery and rapid pathway elucidation in plant specialized metabolism, offering high resolution in gene expression at the cellular level^53,56^. To address the challenge^55^ of performing unbiased annotation of the different cell types in non-model plant species as the majority of medicinal plants are, we developed an automated annotation pipeline using ScType^63^ with a customized marker-gene database based on identification of functional ortholog genes through syntenic analysis.

Our findings reveal a major role of Hyper cells — a previously unidentified type of cells — to function as cellular factories for the biosynthesis *de novo* of hyperforin that combines three biosynthetic modules (MEP, PIBP/polyketide and prenylation) in one cell type. The finding that Hyper cells are adjacent to translucent cavities where hyperforin is stored at high levels in leaves, together with the identification of biosynthetic genes, with our next steps toward identifying regulatory and transport mechanisms will deepen the understanding of specialized metabolism in plants at the single-cell level^79^. Our results not only provide a synthetic-biology approach to produce the hallmark polycyclic scaffold of hyperforin, but also underscore a powerful strategy for accelerating discovery and manipulation of natural-product biosynthesis in medicinal plants that are otherwise not readily amenable to functional characterization in conventional model organisms.

## Materials and Methods

### Plant materials

*Hypericum perforatum* subsp. *chinense* were collected in Enshi Tujia and Miao Autonomous Prefectures in Hubei, China, were delivered and grown under a 16 h:8 h light:dark cycle at 20°C in the CEMPS Phytotron.

### Bulk RNA sequencing

A total of 17 plant tissues, namely untreated leaves, MeJA-treated leaves^20^, young flowers, old flowers, young buds, middle-aged buds, old buds, leaves near buds, young leaves, middle-aged leaves, old leaves, stressed leaves, young stems, old stems, old roots, primary roots, and secondary roots were frozen on liquid nitrogen and homogenized using a tissue lyser (Shanghai Jingxin 40 JXFSTPRP-48). Total RNA was then extracted using the RNAprep Pure Plant Plus Kit (TIANGEN, Cat. #DP441), following the manufacturer’s instructions.

The purity of the extracted RNA was assessed based on the OD_260 nm_/OD_280 nm_ ratio determined by a NanoDrop instrument (Thermo Fisher Scientific, USA) and agarose gel electrophoresis was performed to test for RNA degradation and potential contamination. RNA integrity was evaluated using a 2100 Bioanalyzer (Agilent Technologies, USA).

RNA samples that met quality-control criteria were used as input for preparation of Illumina short-read libraries. The NEBNext Ultra RNA Library Prep Kit (NEB, USA, Cat. #E7530L) was used according to the manufacturer’s recommendations and index codes were incorporated into the libraries to attribute sequences to their respective samples. Qualified libraries were pooled together and subjected to sequencing on the Illumina HiSeq X Ten platform to produce 150-bp pair-end reads. Mixed RNA from 11 different plant tissues (except untreated and MeJA-treated leaves) was prepared to construct continuous long-read (CLR) libraries using the SMRTbell Express Template Prep Kit 2.0 and sequenced on PacBio Sequel IIe to generate a full-length transcriptome.

### Genome sequencing, assembly and annotation

For the long-read sequencing of the St. John’s wort genome, genomic DNA was extracted using a phenol–chloroform method^80^. We constructed Single Molecule, Real-Time (SMRT) sequencing libraries following Pacific Biosciences standard protocols (www.pacb.com) by shearing DNA to a target size of 15–18 kb, followed by damage repair, end repair, blunt-end ligation and size selection to create a large insert SMRTbell library. This library was then sequenced on the PacBio Sequel platform. The assembly of the long reads was carried out using Hifiasm^81^. Additionally, a crosslinking step was employed using 4% v/v formaldehyde for the genomic DNA followed by vacuum treatment. After quenching the crosslinking reaction, samples were digested overnight with MboI. The fragment ends were marked with biotin and the chromatin DNA was re-ligated. The DNA underwent purification via the phenol–chloroform method, and biotin was removed from non-ligated fragment ends. A-tails were then added to the fragment ends and ligated to Illumina paired-end sequencing adapters. The resulting Hi-C sequencing libraries were amplified by PCR and sequenced on the Illumina NovaSeq platform. The data from Hi-C sequencing were used to anchor the long-read assembly into pseudochromosomes.

Repetitive genomic sequences were identified using TRF^82^, RepeatMasker^83^ and LTR_FINDER^84^. Structural annotation was performed using TblastN^85^ and GeneWise^86^ for homolog prediction and tools Augustus^87^, Geneid^88^, and Genescan^89^ for *ab initio* gene prediction. Integration of RNA-seq data facilitated accurate gene-model prediction. Functional annotation involved aligning proteins to the Swiss-Prot database and domains to InterProScan. The prediction of non-coding RNAs, including tRNAs, rRNAs, miRNAs and snRNAs, was achieved using tRNAscan-SE^90^ and alignment with the Rfam database. Genomic features were presented by different tracks in a CIRCOS plot^91^.

### Reference-genome-based read mapping

To acquire high-quality, clean RNA-seq reads, the raw data were trimmed using Trimmomatic v0.36^92^ with the following parameters: HEADCROP: 13; LEADING: 3; TRAILING: 3; SLIDINGWINDOW: 4:15; and MINLEN: 36. Clean data were then aligned to the genome assembly using hisat2^93^ with default settings. Fragments Per Kilobase of transcript per Million mapped reads (FPKM) values were then generated through the cuffdiff command within the Cufflinks suite^94^.

### Preparation of St. John’s Wort protoplasts for single-cell RNA-seq

Fresh St. John’s Wort leaves and buds from 1 year old plants grown in the CEMPS Phytotron were collected and thoroughly washed with distilled water to remove any surface contaminants. Anthers were removed from bud samples because of their negative influence on protoplast purity, leaving only the petals and ovary. Tissues were then finely chopped and placed in a 10 cm culture dish. 10 mL of digestion solution (pH = 5.7) containing cellulase (1.25% w/v), pectinase (0.3% w/v), *D*-mannitol (0.4 M), MES (20 mM) and KCl (20 mM) was added to the culture dish containing the macerated tissues and incubated at room temperature (25 °C) with shaking at 80 rpm for 2 h. The digested solution was centrifuged at room temperature (500 *g*) for 10 min to separate the protoplast- containing pellet from the tissue debris. The resulting protoplast pellet was carefully re-suspended in 500 μL washing buffer consisting of 0.4 mM mannitol, 20 mM MES pH 5.7, 20 mM KCl, 10 mM CaCl_2_, and 0.1% (w/v) BSA. The protoplast suspension was filtered through a 40-µm cell sieve to remove any remaining tissue debris and obtain a clarified protoplast solution, followed by centrifugation at room temperature (200 *g*) for 6 min to further remove debris and obtain a purified protoplast pellet. The pellet was gently resuspended in 40 μL fresh washing buffer to obtain a concentrated protoplast suspension suitable for subsequent experiments. The viability of the isolated protoplasts was evaluated using trypan blue staining.

### Single-cell (sc) RNA sequencing (scRNA-seq)

The library construction process used the Chromium Single Cell 3ʹ Library & Single Cell 3ʹ v3 Gel Beads kit. To generate single-cell GEMs (gel beads in emulsion), a suspension of protoplasts was loaded onto the automated Chromium Controller instrument. Ensuring the integrity of each GEM, reverse transcription occurred within them, resulting in cDNA that carried a unique cell barcode. Following cDNA synthesis, GEMs were broken to release the cDNA from individual cells. Liberated cDNA was then pooled and amplified using PCR. After amplification, the cDNA was fragmented and sequencing adapters were attached to construct libraries. Using the Illumina HiSeq X Ten platform, paired-end short reads of 150 bp were generated.

### scRNA-seq data analysis

Raw sequencing data were preprocessed to transform it into a structured format by Cellranger^95^. Initial steps in Seurat (ref) involved filtering the dataset to exclude cells with aberrantly high gene counts (> 3,000), potentially indicative of artifacts or multiplets. This was followed by normalization to adjust for variations in sequencing depth across individual cells using 10,000 as scale factor. To focus the analysis on genes most likely to reveal biologically significant variation, a feature-selection process based on the top 2,000 variable genes was used. PCA dimensionality reduction was then applied to the data to distill the high-dimensional gene-expression profiles. The selection of dimensions for this reduction was determined through jackstraw and elbow plot^58^. The presence of potential doublets was assessed and removed by DoubletFinder^96^. Clustering algorithms were then used to identify distinct cell populations within the dataset. The resolution of clustering was fine-tuned by clustree^97^ ranging from 0.1 to 1. For visual representation, the cell populations were projected by UMAP reduction. Marker genes were identified as genes predominantly expressed in each cell cluster. UCell^98^ was used to score gene signatures based on the gene data matrix and MetaNeighbor^73,74^ was used to quantify cell-type similarity between two datasets using neighbor voting.

We adapted a customized cell-type annotation approach using the ScType pipeline ^63^, due to the limitation of the standard ScType marker gene database being specific to human and mouse. We compiled an extensive marker gene dataset for *A. thaliana* from PlantscRNAdb ^59^and mapped these genes to their syntenic counterparts in *H. perforatum*. This tailored database enabled to perform automated and unbiased cell-type classification. A schematic of this annotation is detailed in Fig. S9.

### Phylogenetic analysis of prenyltransferases

The amino-acid sequences of characterized prenyltransferases were aligned with sequences of genes obtained from synteny analysis using MUSCLE^99^. JTT+I+G was selected as the best-fit model of amino-acid replacement using prottest^100^ based on alignment results. The maximum-likelihood method was then used to infer the phylogenetic tree by RaxML-ng^101^ with 1,000 bootstrap values.

### Gene isolation and plasmid construction

Total RNA was extracted from the aerial parts of *H. perforatum* and 5 μg was transcribed into cDNA using the SuperScriptTM IV First-Strand Synthesis system (ThermoFisher, Invitrogen). Candidate gene ORFs were PCR-amplified (Table S17) and cloned into various pESC vectors (pESC-LEU for *HpCCL2* and *HpPKS2*, pESC-HIS for *HpPT1* and *HpPT2*, pESC-URA for *HpPT3* and *HpPT4*), and pEAQ-HT (for *N. benthamiana* expression), PA7-YFP (yellow fluorescent protein was fused to the C-terminus of prenyltransferase signal peptides) using the ClonExpress II One Step Cloning Kit (Vazyme). Restriction enzymes and overhangs are detailed in Table Y. Following transformation into *E. coli* DH5α (DE3), colonies were selected on antibiotic-supplemented LB agar. Positive colonies were confirmed through colony PCR using primers listed in Table S17 and plasmids were isolated from cultures grown overnight at 37°C for verification by sequencing.

### Expression and enzyme activity of prenyltransferases in yeast

The *Saccharomyces cerevisiae* DD104 strain (engineered for isoprenoid production)^102^ was transformed using lithium acetate to introduce combinations of *HpCCL2*, *HpPKS2* and *PT* candidate genes inserted into pESC vectors (Agilent). Three days after transformation, colonies were grown in 5 mL dextrose dropout medium for 2 d. Following this, cells were harvested and re-suspended in 15 mL of dropout medium containing 2% (w/v) d-galactose for 3 d. After induction, cultures were extracted using an equal volume of ethyl acetate (EtOAc). The organic layer obtained was separated and dried under reduced pressure. The dried extract was subsequently resuspended in 400 µL MeOH followed by filtration using a 0.22-μm pore-size syringe filter.

### Transient expression in *Nicotiana benthamiana*

For *Agrobacterium*-mediated transient expression in *N. benthamiana*, pEAQ-HT plasmids (Plant Bioscience Limited, Norwich, UK)^103^ harboring genes of interest were transformed into *Agrobacterium tumefaciens* GV3101 with selection. PCR-positive colonies were cultured overnight in 15 mL LB medium with relevant selection. Cells were centrifuged and re-suspended in MMA buffer (10 mM MgCl_2_, 10 mM MES, 150 μM acetosyringone), adjusted to OD_600 nm_ = 0.2 and kept in the dark for 2 h. Leaves of 4–6 week-old *N. benthamiana* were infiltrated with the indicated combinations of each suspension. After 3 d, plants were supplemented with 2mM valine solution, and 2 d later, leaves were collected, flash-frozen and stored at –80°C. Frozen leaves were then lyophilized and ground into a fine powder then extracted in 10 mL methanol. The extraction was concentrated under vacuum, re-suspended in 500 µL methanol, filtered through a 0.22-µm organic syringe filter and stored at –20°C.

### LC–MS method for metabolite analysis

Metabolites from co-expression experiments in yeast and tobacco samples were subjected to analysis using a Q Exactive system, integrated with a Dionex Ultimate 3000 UHPLC system. Three microliters (3 µL) of each processed sample was injected into a UPLC (Kinetex 2.6 µm C18 100 Å, measuring 100 x 2.1 mm, Phenomenex). This was carried out at a flow rate of 0.35 mL per minute. The gradient with solvent A (2 mM NH_4_Ac) and solvent B (MeCN) from 0 to 0.5 min with maintained 10% B, followed by a linear increase from 10% to 98% B between 0.5–11 min. This was sustained at 98% B from 11–13 min. Mass spectra acquisition was performed in the positive mode under the following parameters: a spray voltage of 3,500 V, a capillary temperature of 320°C, a sheath gas of 40 and an aux gas of 10.

### *In situ* RNA hybridization

RNA *in situ* hybridization was conducted using an optimized protocol^104^. Young *H. perforatum* leaves were fixed in FAA solution (3.7% v/v paraformaldehyde, 5% v/v acetic acid and 50% v/v ethanol), dehydrated by Leica HistoCore PEARL, and embedded in paraffin by Leica HistoCore Arcadia H. Thin sections (8 μm) were prepared using the HistoCore MULTICUT. The sections underwent toluidine blue staining for morphological examination. Additionally, a schematic representation of the leaf was generated by ggplantmap^105^ for detailed visualization. Sense and antisense probes were synthesized through T_7_-promoter-based *in vitro* transcription, incorporating digoxigenin-labeled UTP for subsequent detection. Hybridization was performed overnight at room temperature allowing for optimal probe–target annealing. Post-hybridization washes were conducted to remove excess probe using a series of saline-sodium citrate (SSC) buffers of decreasing concentration. The hybridized probes were then detected using an anti-digoxigenin antibody conjugated to alkaline phosphatase, with NBT/BCIP as the chromogenic substrate. The specificity of the hybridization signal was confirmed by comparing the results with sense probe controls. The hybridized sections were visualized using an Olympus BX53 microscope.

### Subcellular localization of prenyltransferases

*Arabidopsis thaliana* protoplasts were prepared from Columbia-0 ecotype and transformed with PA7-YFP plasmids as described previously^20^. Yellow fluorescent protein (YFP) was fused to the C-terminus of predicted prenyltransferase signal peptides alone, giving rise to HpPT1–YFP (comprising amino acids 1–76), HpPT2–YFP (1–75), HpPT3–YFP (1–71) and HpPT4–YFP (1–79). The images were captured with a Zeiss LSM880 confocal microscope.

### Chemical synthesis of prenylated intermediates

For translucin-I synthesis, PIBP (2.55 mmol) and geraniol (2.55 mmol) were dissolved in anhydrous CH_2_Cl_2_ (5 mL), followed by the addition of BF_3_·Et_2_O (0.25 mmol) at 35°C. The reaction mixture was stirred for 1 h at the same temperature. Post-reaction, it was quenched with saturated NaHCO_3_ solution (10 mL) and extracted with CH_2_Cl_2_ (3 × 20 mL). The dried organic phases (over Na_2_SO_4_) were then purified using silica gel column chromatography with EtOAc−hexane to yield translucin I^106^.

For the synthesis of translucin II and III, translucin I (0.2 mmol) was dissolved in H_2_O (2 mL) and THF (1 mL) under nitrogen stream at 0°C. KOH (0.6 mmol) was added, followed by prenyl bromide (0.8 mmol). After stirring for 1 h at 0°C, the reaction was quenched with saturated NaHCO_3_ solution (10 mL) and extracted with CH_2_Cl_2_ (3 × 10 mL). The combined organic phases were dried over Na_2_SO_4_ and purified using silica gel column chromatography with EtOAc−hexane, resulting in translucin II and III^107^.

## Acknowledgments

We would like to thank the staff and management of the CEMPS Core Facility Center for the support in metabolomics (LC-MS, NMR) and microscopy services, as well as the personnel at the CEMPS Phytotron facilities. We also thank Prof. Bin Han and Dr Tao Huang for the access to computational servers for the bioinformatic analysis at the National Centre for Gene Research. We thank Wenjie Liang and Haojie Li for the help of microscope and Kexin Ji and Jiang Wang for the help of heterologous expression in tobacco. We acknowledge Prof. George Lomonossoff (John Innes Centre) for providing us with the pEAQ-HT plasmid. We acknowledge Prof. Guodong Wang (IGDB) for providing us with the DD104 yeast strain and the hops genes *HlPT1L*, *HlPT2*.

The work was financially supported by the National Natural Sciences Foundation of China, Research Fund for International Excellent Young Scientists (RFIS-II), grant 32150610477; Strategic Priority Research Program of the Chinese Academy of Sciences grant XDB27020204; Chinese Academy of Sciences, International Partnership Program of CAS grant 153D31KYSB20160074; and National Key Laboratory of Plant Molecular Genetics Special Fund. The CAS PIFI Fellowship and the China Postdoctoral Science Foundation for Postdoctoral International Exchange Program Fellowship provided fellowship support to Dr Ana Luisa Malaco Morotti.

