## Supplementary material for "Single-cell RNA-seq based elucidation of the antidepressant hyperforin biosynthesis *de novo* in St. John’s wort": SI

**The file includes**:
Figures S1 to S31.

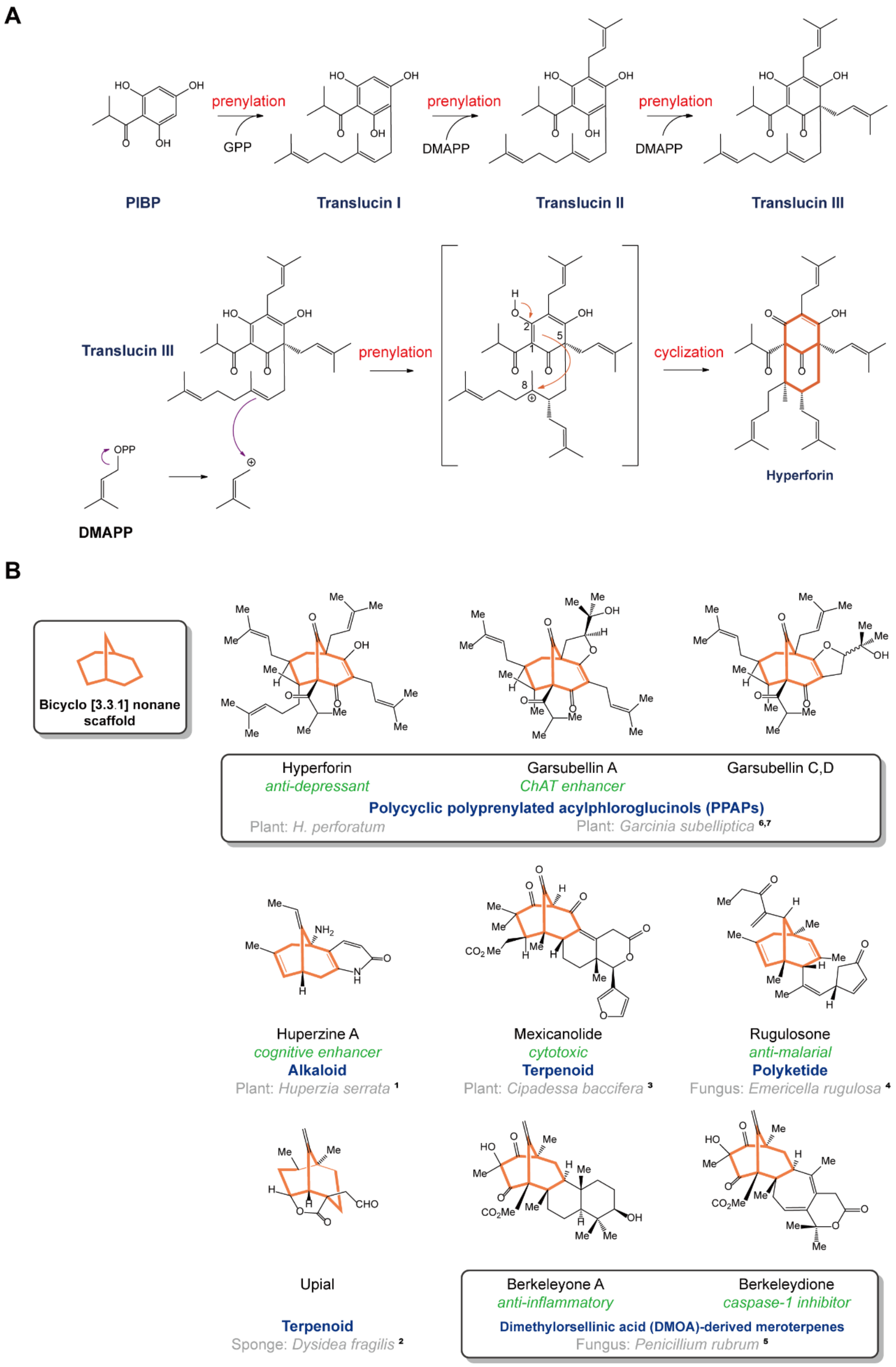


**Figure S1. Hyperforin biosynthesis and the formation of bicyclo[3.3.1]nonane scaffold (highlighted in orange).**

**A.** Hyperforin biosynthetic pathway and tentative mechanism of bicyclo[3.3.1]nonane scaffold creation through the formation of a carbocation intermediate. A similar hypothesis has been proposed by Arigoni[1], Dewick[2] and Walsh[3].

**B.** The occurrence of the bicyclo[3.3.1]nonane scaffold in bioactive natural products across plants, fungi and animals[4]. The key bicyclo[3.3.1]nonane scaffolds are highlighted in orange, bioactivities in green and host organism in gray.

**1**, Jia-Sen Liu *et al*. The structures of huperzine A and B, two new alkaloids exhibiting marked anticholinesterase activity. *Canadian Journal of Chemistry*. 64(4): 837-839 (1986).

**2**, Gary Schulte *et al*. *The Journal of Organic Chemistry* 1980 45 (3), 552-554.

**3**, Cao DH *et al*. Mexicanolide-type limonoids from the twigs and leaves of *Cipadessa baccifera*. *Phytochemistry*. Epub Jun 26 (2020).

**4**, Moosophon P *et al*. Prenylxanthones and a bicyclo[3.3.1]nona-2,6-diene derivative from the fungus *Emericella rugulosa*. *J Nat Prod*. 2009 Aug;72(8):1442-6.

**5**, Stierle DB *et al*. Berkeleydione and berkeleytrione, new bioactive metabolites from an acid mine organism. *Org Lett*. 2004 Mar 18;6(6):1049-52.

**6**, Fukuyama Y *et al*. Garsubellin A, a novel polyprenylated phloroglucin derivative, increasing choline acetyltransferase (ChAT) activity in postnatal rat septal neuron cultures. *Chem Pharm Bull (Tokyo)*. 1997 May;45(5):947-9. Doi: 10.1248/cpb.45.947. PMID: 9178529.

**7**, Fukuyama Y *et al*. Garsubellins, polyisoprenylated phloroglucinol derivatives from *Garcinia subelliptica*, *Phytochemistry*, Volume 49, Issue 3, 1998, Pages 853-857, ISSN 0031-9422.

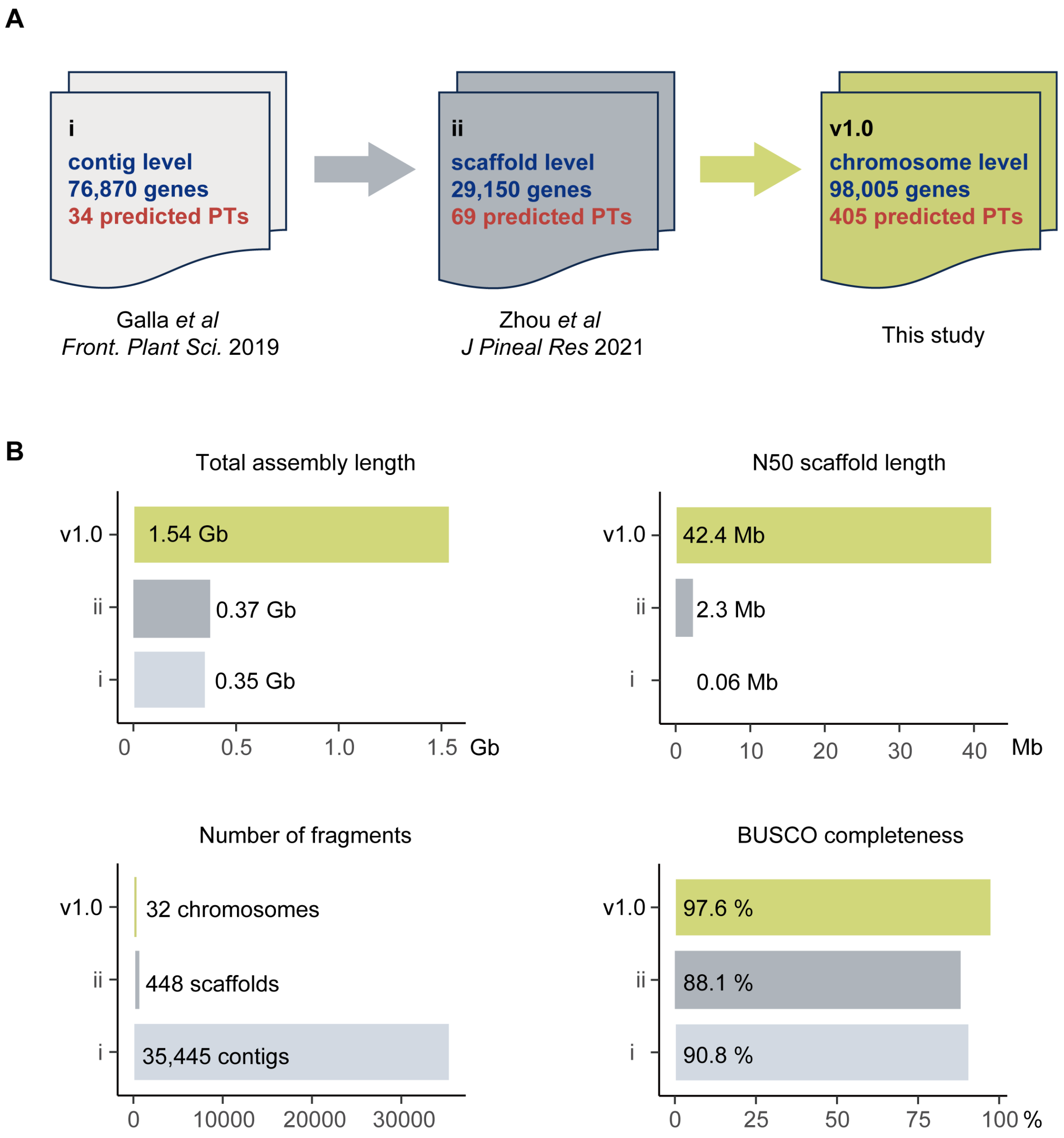


**Figure S2. Advances of the current reference tetraploid *H. perforatum* genome published in this study compared with previous versions.**

**A.** Comparison of continuity, total-gene numbers and numbers of predicted prenyltransferases across three *H. perforatum* genome assemblies.

**B.** Comparison of total-assembly length, N50-scaffold length, number of fragments and BUSCO completeness across three *H. perforatum* genome assemblies.

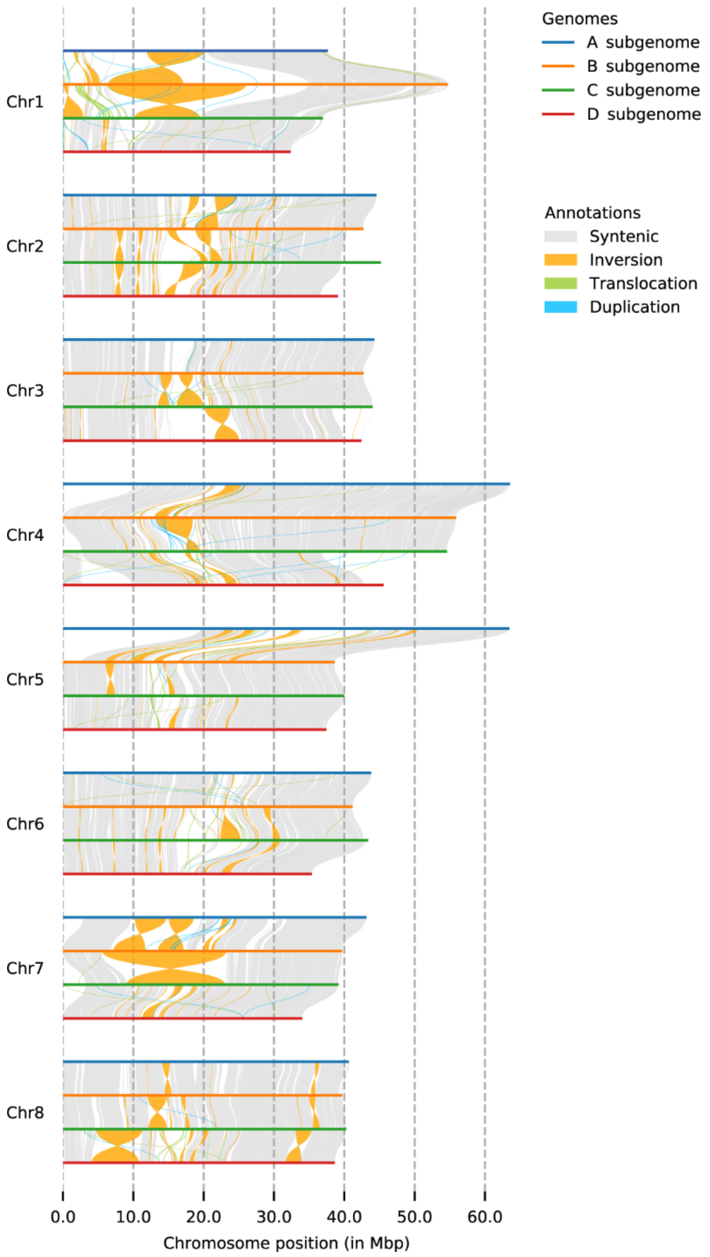


**Figure S3. Synteny and genomic-segment re-arrangements among homologous *H. perforatum* chromosomes .**

Each subgenome A, B, C, and D is depicted by different colored lines (blue, orange, green, and red, respectively). The different color correlation lines (gray, gold, pale green, pale blue) between different subgenomes refer to the observed structural arrangements (syntenic, inversion, translocation, and duplication, respectively).

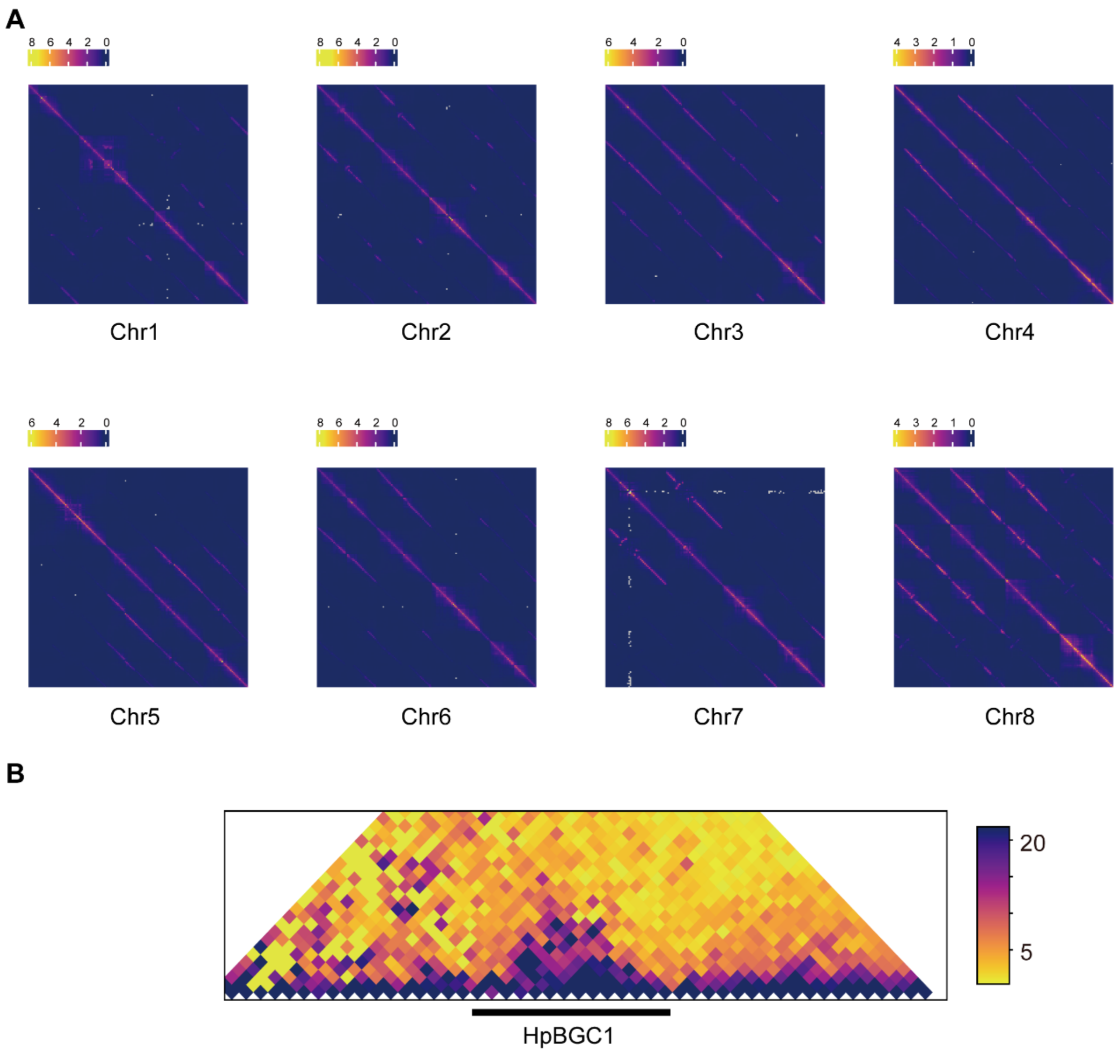


**Figure S4. Hi-C interaction frequencies in the *H. perforatum* genome.**

**A**, Hi-C interaction frequencies within the set of four homologous chromosomes. The color bar indicates contact frequencies from low (purple) to high (yellow).

**B**, Identification of the *BGC1* biosynthetic gene cluster within a topologically associating domain (TAD) by TAD calling. The black line delineates the boundaries of the cluster within the genomic architecture. The color gradient represents interaction frequencies within the TAD, ranging from low (yellow) to high (purple).

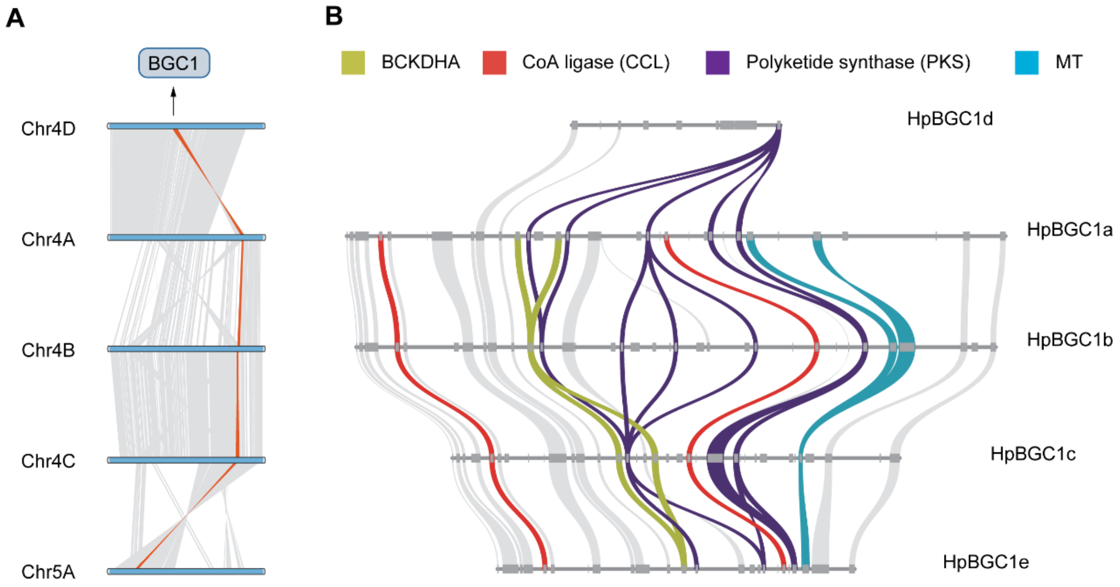


**Figure S5. Syntenic relationship of *Biosynthetic Gene Cluster 1* (*HpBGC1*) in the genome of *H. perforatum*.**

**A**, Macrosynteny of *HpBGC1* across Chr5A and homologous chromosomes Chr4A, Chr4B, Chr4C and Chr4D. Gray wedges in the background indicate major syntenic blocks spanning ≥ 100 genes between the genomes, with syntenic regions harboring *HpBGC1* highlighted in orange.

**B**, Microsynteny of *HpBGC1*, yellow–green ribbons show the *BCKDHAs*; red ribbons show the CoA-ligases (*CCLs*); purple ribbons show genes encoding polyketide synthases (*PKSs*); and blue ribbons show the methyltransferases (*MTs*) that act on the polyketide PIBP.

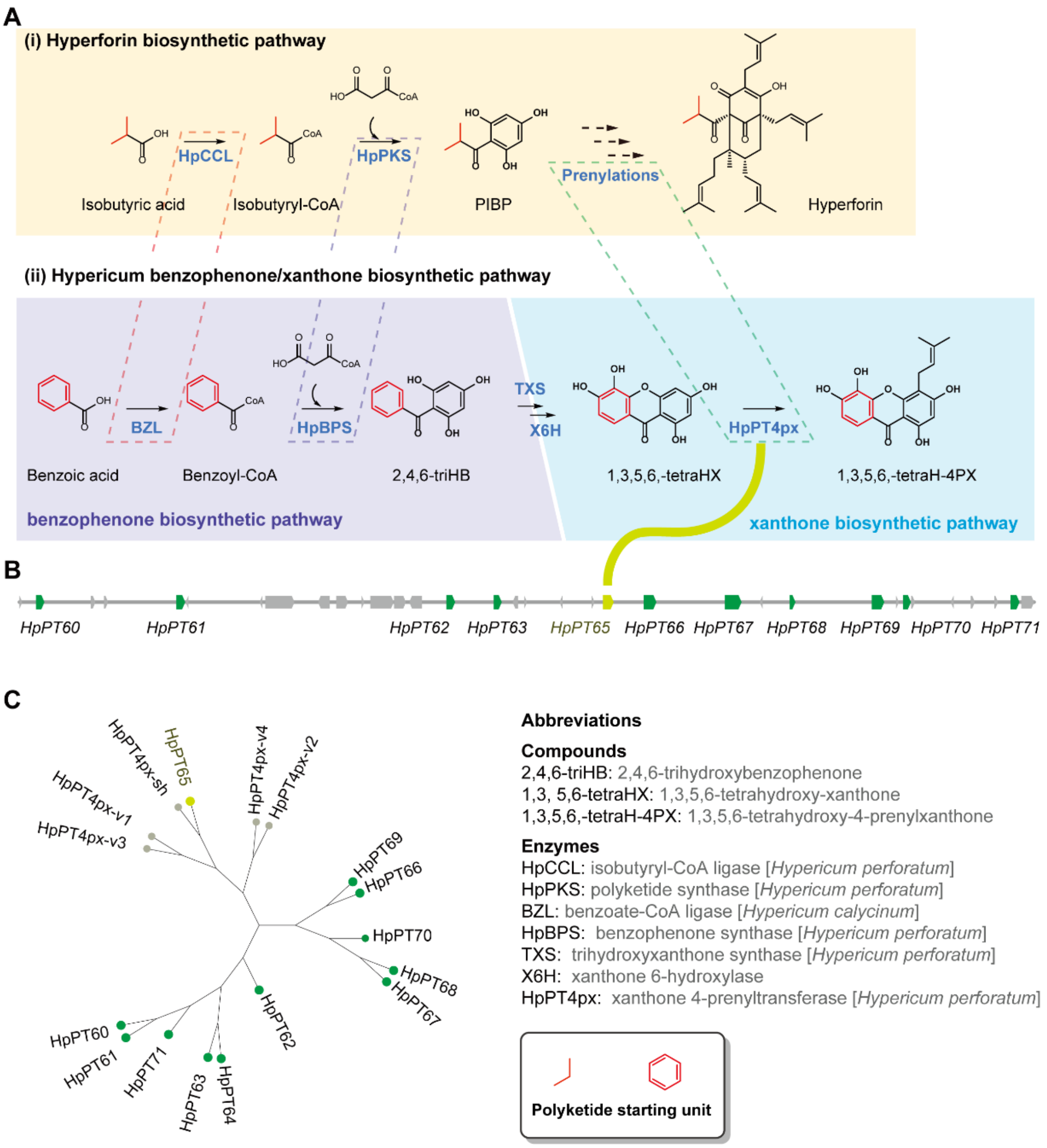


**Figure S6. Identification of a genomic region enriched with prenyltransferase genes in *H. perforatum*.**

**A**, The analogy in the biosynthetic pathways of hyperforin and benzophenones/xanthones, particularly focusing on the initial two steps involving CoA-ligase and polyketide synthase that establish the polyketide scaffold, followed by subsequent prenylation processes.

**B**, Depiction of the *Hypericum perforatum* genomic region highly enriched with the 11 PT genes, with the HpPT65 to be identified as the specific HpPT4px[5] variant in this region.

**C**, Phylogenetic analysis of characterized HpPT4px variants in relation to PTs within this region, based on amino acid sequence alignments.

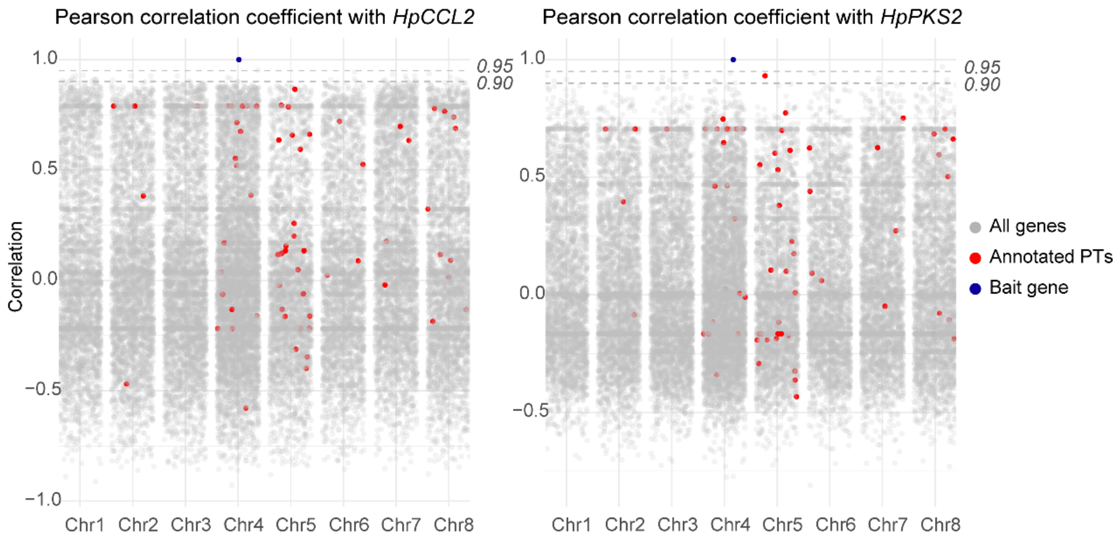


**Figure S7. Co-expression analysis illustrating Pearson correlation coefficients[6] with characterized genes in hyperforin biosynthesis.**

Gray dots represent all analyzed genes, characterized genes in a previous study, *HpCCL2* and *HpPKS2*, are distinguished in blue and genes annotated as PTs are marked in red. Thresholds of significance at Pearson correlation coefficients of 0.90 and 0.95 are distinctly highlighted. The only prenyltransferase gene with Pearson correlation coefficients over 0.90 was *HpPT12,* which was inactive towards PIBP and translucins I–III.

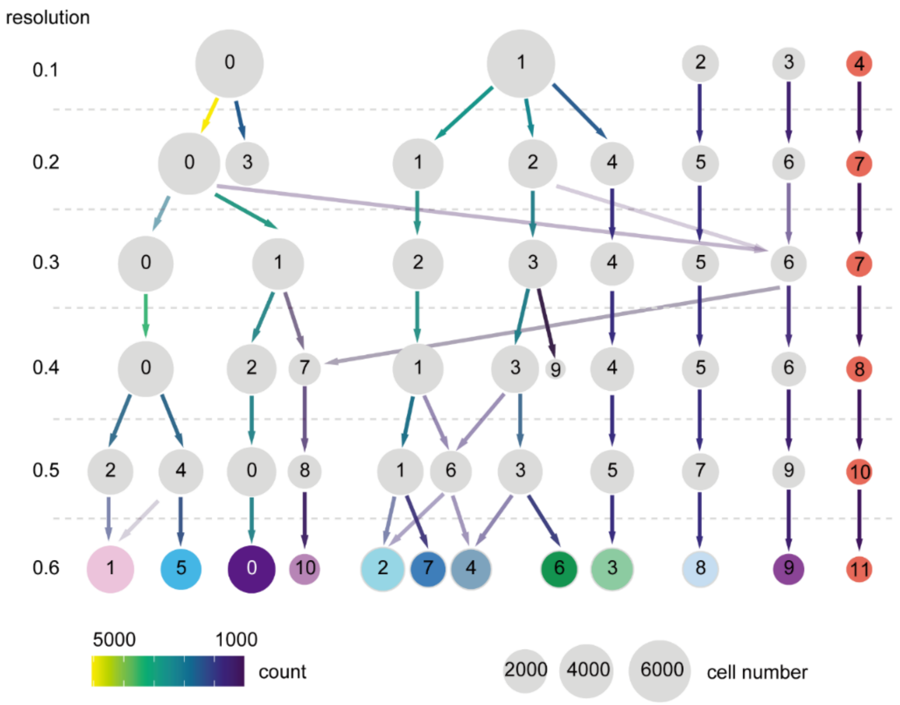


**Figure S8. Clustree[7] visualization of leaf single-cell RNA-seq data across various resolutions.**

The plot illustrates the clustering dynamics of single-cell RNA-seq data at incremental resolutions ranging from 0.1 to 0.6, with each step increasing by 0.1. Cluster 11 (Hyper cells), identified at the resolution of 0.6, consistently emerges as an independent cell cluster across all examined resolutions.

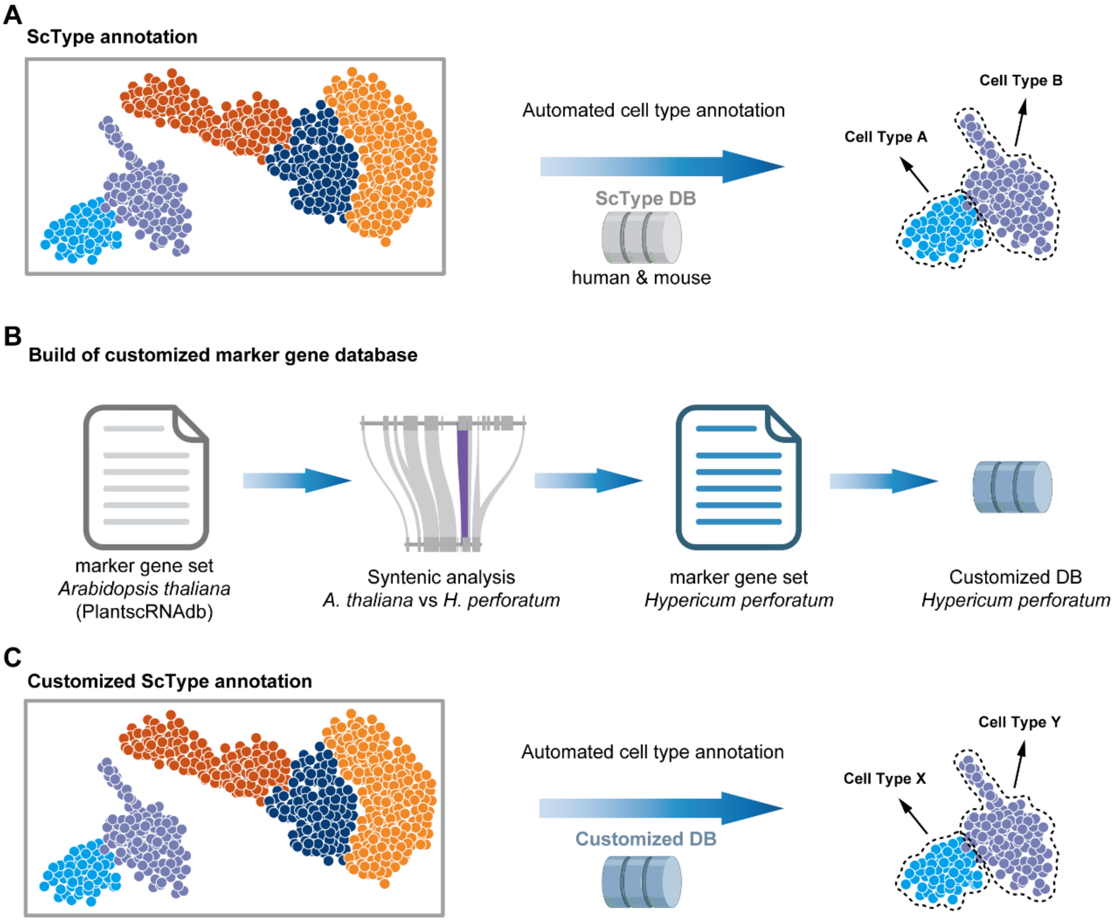


**Figure S9. Overview of the customized single-cell annotation pipeline for *H. perforatum* single-cell RNA atlases**.

**A**, Standard ScType[8] annotation using in-built databases primarily based on human and mouse gene sets.

**B**, Expansion of the annotation process by collecting marker-gene sets from *Arabidopsis thaliana* via PlantscRNAdb[9]. The list is then subjected to synteny analysis with *H. perforatum* genes to derive *H. perforatum*-specific orthologous marker genes, thus constructing a tailored database.

**C**, Implementation of a customized ScType annotation using the newly developed *H. perforatum*-specific database.

A

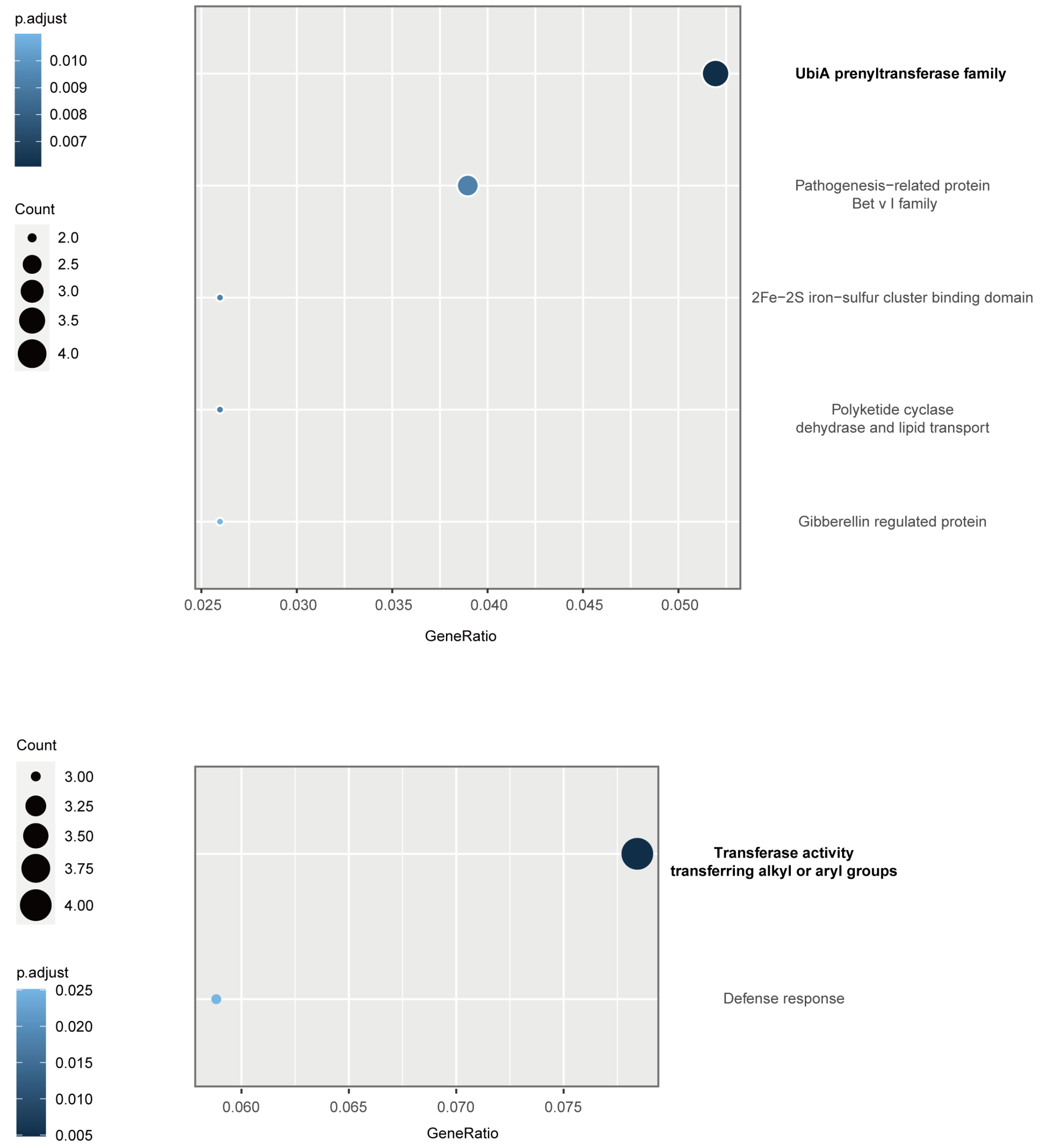


**Figure S10. Enrichment analysis of Hyper cells in the *H. perforatum* leaf atlas.**

**A**, Pfam domain-enrichment analysis reveals that the most significantly enriched term is the 'UbiA prenyltransferase family'.

**B**, Gene Ontology (GO) term enrichment analysis identifies 'transferase activity' as the top-ranked term.

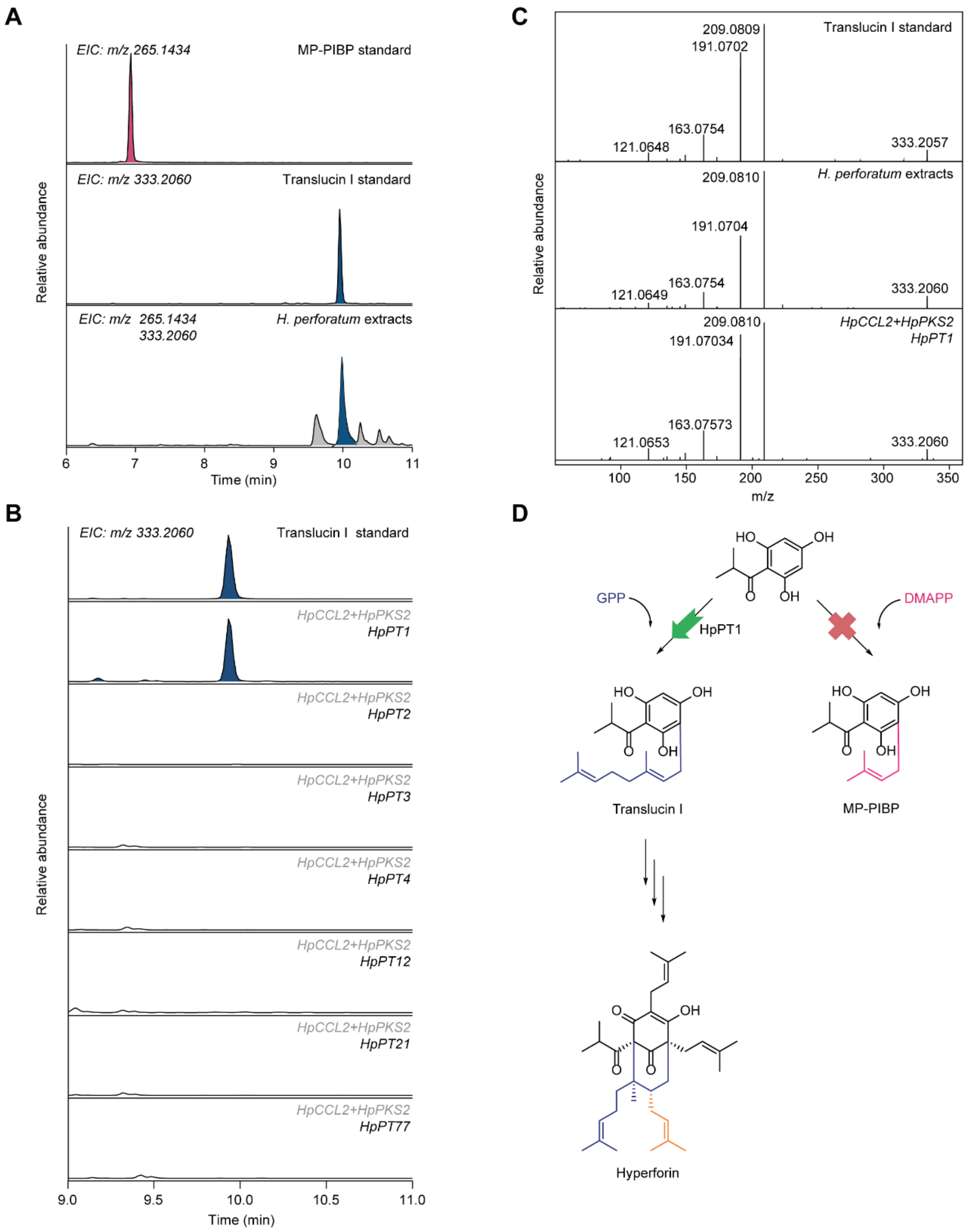


**Figure S11. HpPT1-catalyzed geranylation of PIBP to form translucin I in *H. perforatum*.**

**A**, LC–MS analysis of *H. perforatum* extracts at selected extracted ion chromatograms (EICs) showing the absence of monoprenylated phloroisobutyrophenone MP-PIBP but the presence of translucin I.

**B**, LC–MS extracted ion chromatograms EIC at m/z 333.2060 depicting that only the co-expression of *HpPT1* with *HpCCL2* and *HpPKS2* results in a distinct new peak, corresponding to the standard of translucin I.

**C**, Comparative MS/MS fragmentation analysis of translucin I standard, corresponding to compounds in *H. perforatum* extracts and the product from the *HpPT1* experiment, confirming the identity of the geranylated product.
**D**, The absence of MP-PIBP and presence of translucin I in extracts coupled with the demonstrated activity of HpPT1 in converting PIBP to translucin I and not MP-PIBP, suggests that HpPT1 selectively uses only geranyl-diphosphate (GPP) as a prenyl donor.

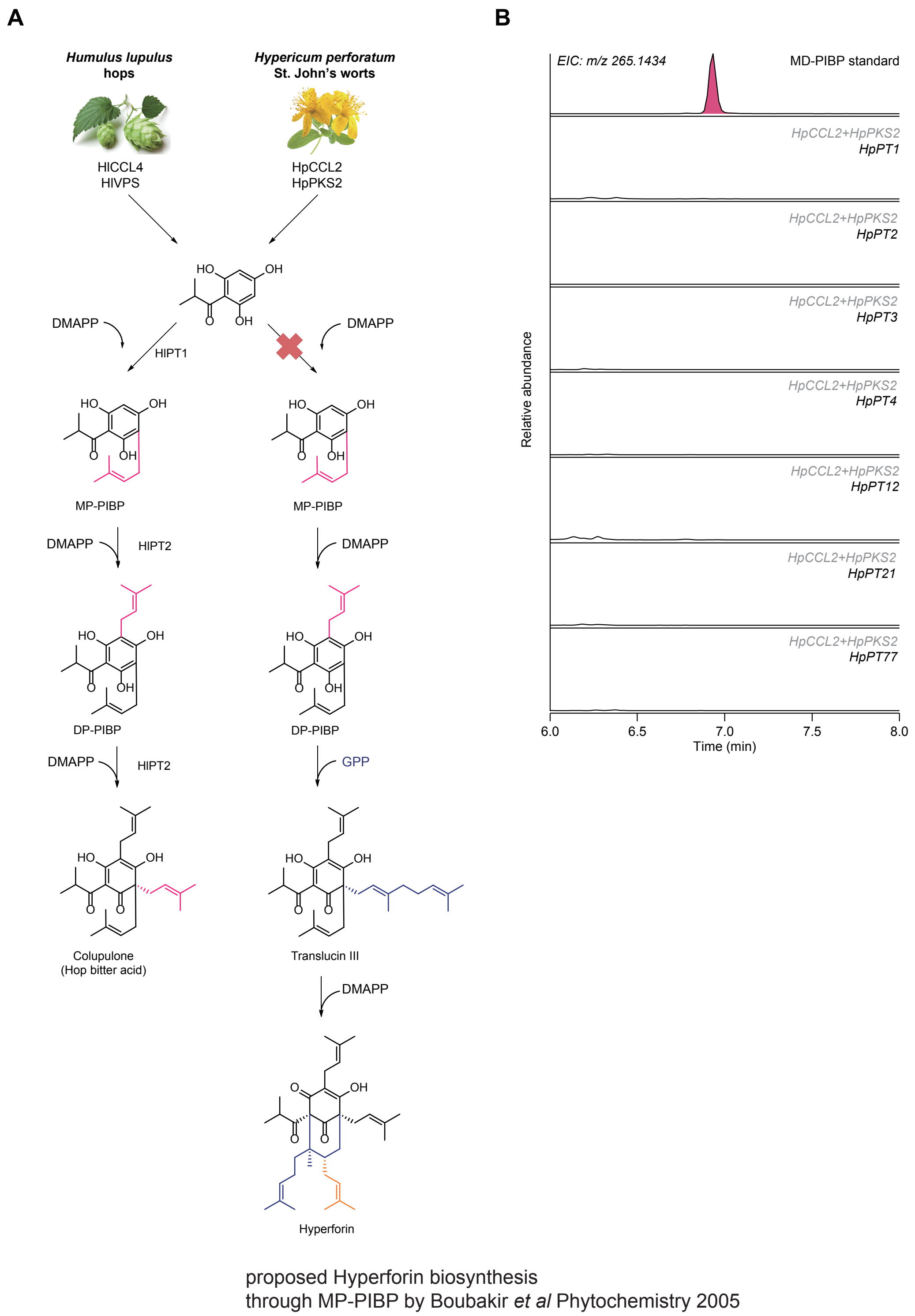


**Figure S12. Refuting the proposed role of MP-PIBP as an intermediate in hyperforin biosynthesis.**

**A**, Illustration of a previously proposed biosynthetic pathway for hyperforin in analogy with the biosynthesis of the hop bitter-acid colupulone suggesting MP-PIBP is an intermediate[10]. Due to chemical similarities, it was supported by biochemical homology between the biosynthesis *de novo* of hyperforin and hop bitter acids. The biosynthesis *de novo* of hyperforin and colupulone have evolved by convergence[11] and MP-PIBP is not a precursor in hyperforin biosynthesis[10].

**B**, LC–MS chromatogram at EIC m/z 265.1434 of the MP-PIBP synthetic standard in comparison with the *in vivo* co-expression experiments involving prenyltransferase candidate genes with *HpCCL2* and *HpPKS2*. The absence of MP-PIBP peaks, when compared against the synthetic standard, suggests that MP-PIBP is not catalyzed by any of the tested prenyltransferases.

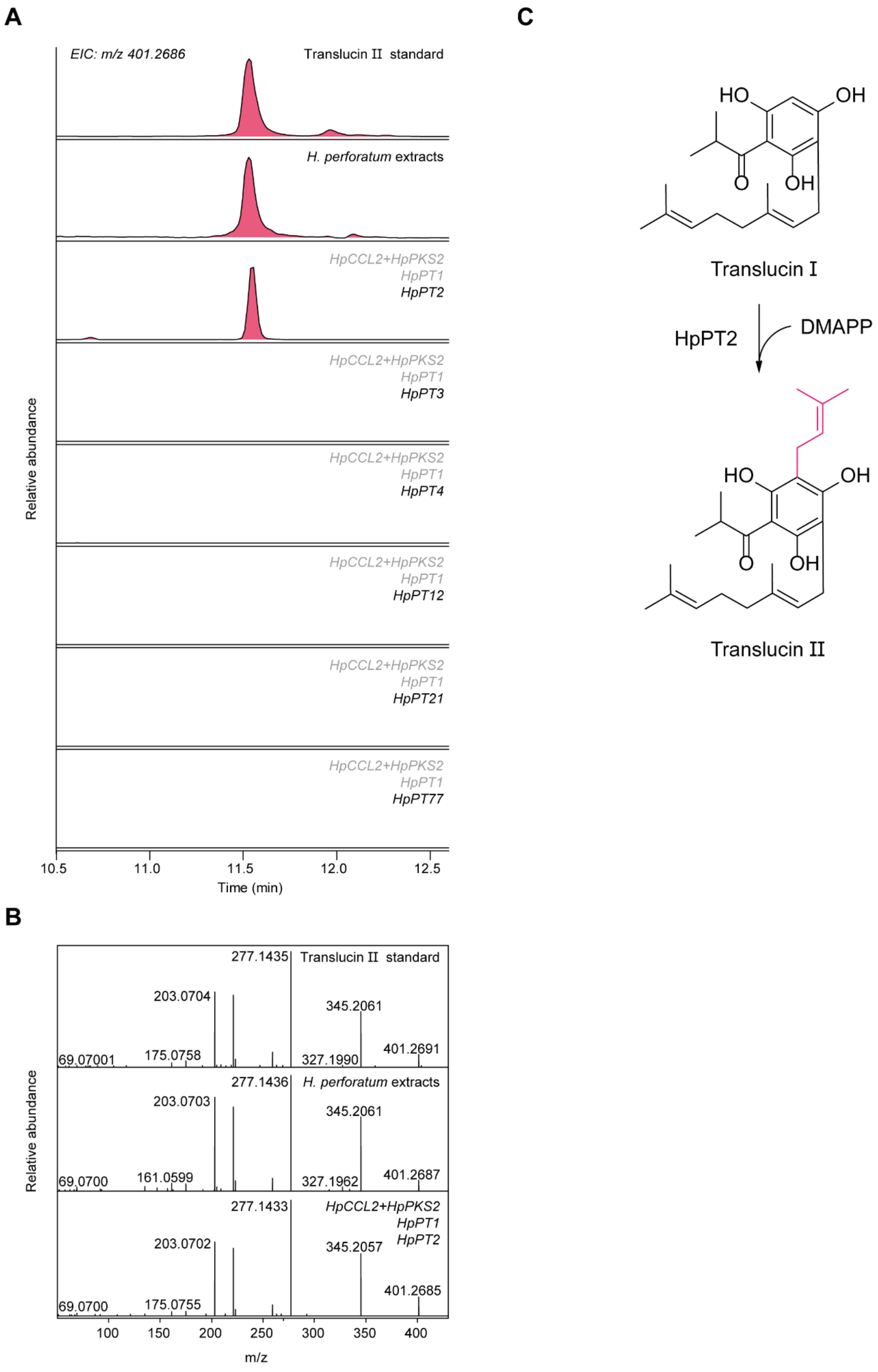


**Figure S13. HpPT2-catalyzed prenylation of translucin I to translucin II in *H. perforatum*.**

**A**, LC–MS extracted ion chromatograms EIC at m/z 401.2686 of the translucin II synthetic standard, a *H. perforatum* plant extract and the ethylacetate extracts from yeast combinatory co-expression of candidate prenyltransferases and upstream metabolic genes for biosynthesis of translucin I. Only the co-expression of *HpPT2* with *HpCCL2*, *HpPKS2*, and *HpPT1* results in a distinct new peak, corresponding to the standard of translucin II.

**B**, Comparative MS/MS fragmentation analysis of the translucin II standard, the corresponding compound in an *H. perforatum* extract, and the product from the HpPT2-characterization experiment confirming the identity of the prenylated product as translucin II.

**C**, Schematic of the enzymatic transformation from translucin I to translucin II catalyzed by HpPT2.

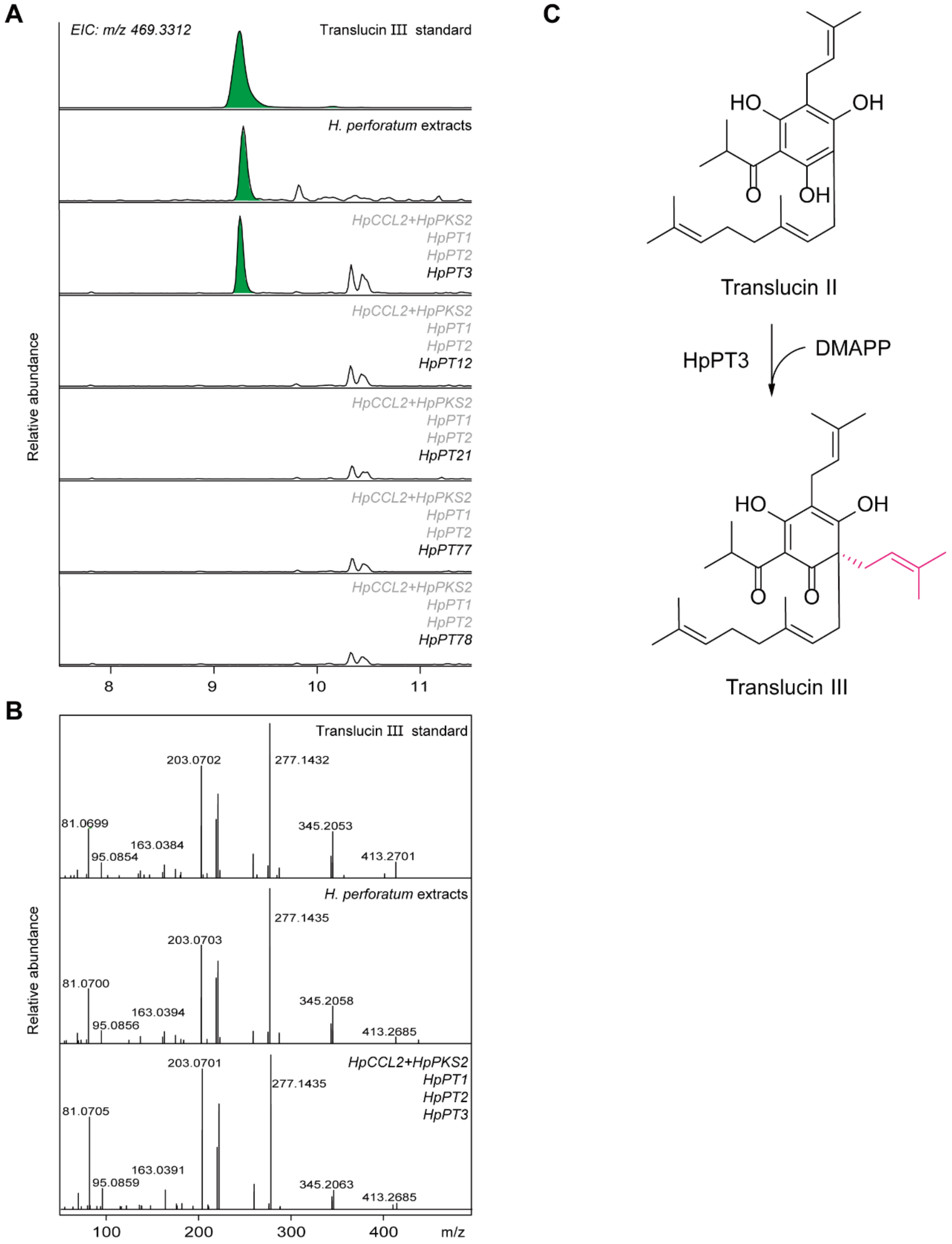


**Figure S14. HpPT3-catalyzed prenylation of translucin II to translucin III in *H. perforatum*.**

**A**, LC–MS extracted ion chromatograms EIC at m/z 469.3312 of the translucin III synthetic standard, an *H. perforatum* plant extract and the ethylacetate extracts from yeast combinatory co-expression of candidate prenyltransferases and upstream metabolic genes for biosynthesis of translucin II. Only the co-expression of *HpPT3* with *HpCCL2*, *HpPKS2*, *HpPT1* and *HpPT2* results in a distinct new peak, corresponding to the standard of translucin III.

**B**, Comparative MS/MS fragmentation analysis of the translucin III standard, the corresponding compound in an *H. perforatum* extract and the product from the HpPT3-characterization experiment confirming the identity of the prenylated product as translucin III.

**C**, Schematic of the enzymatic prenylation of translucin II to translucin III catalyzed by HpPT3 and the subsequent dearomatization of the polyketide PIBP ring.

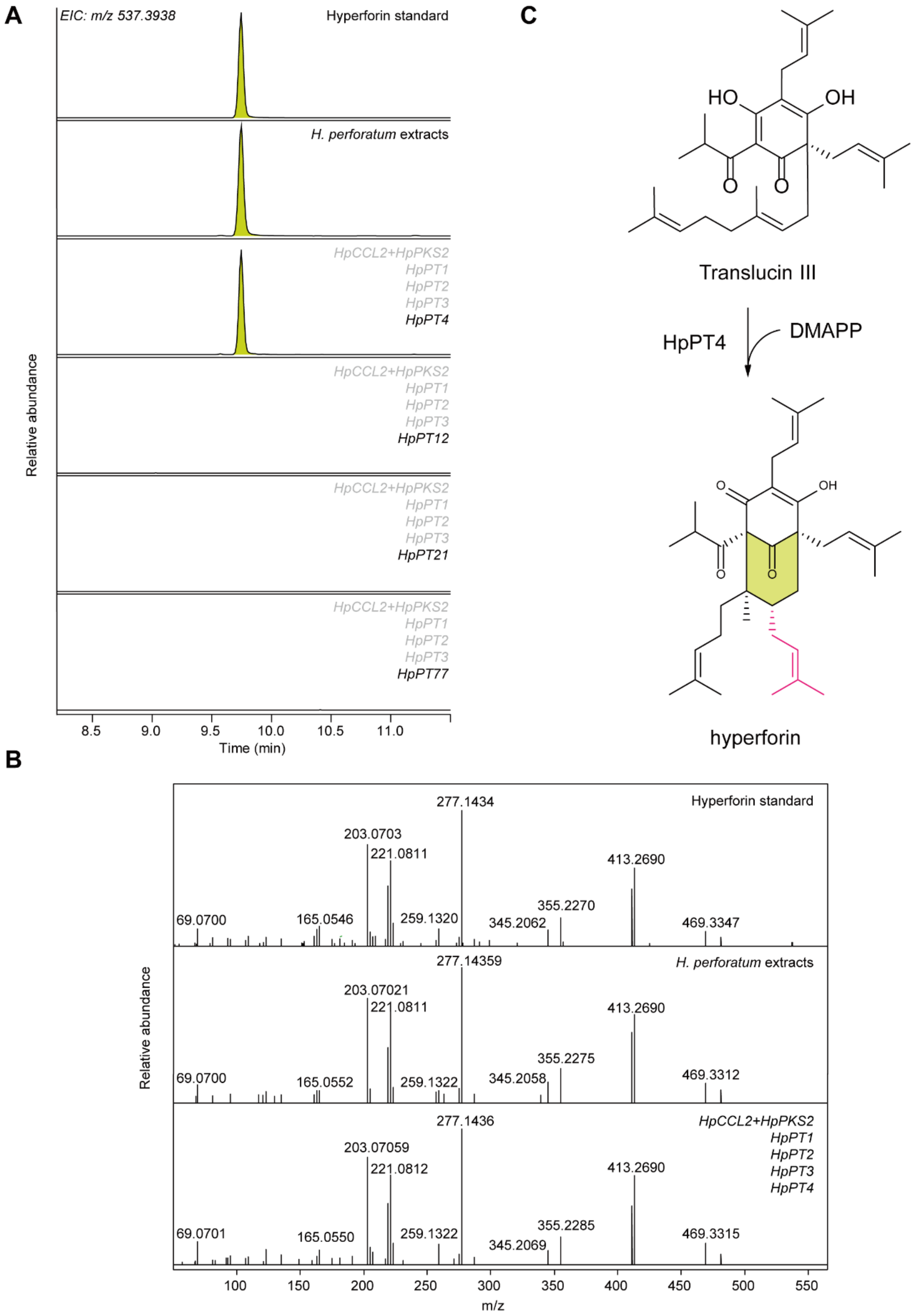


**Figure S15. HpPT4-catalyzed prenylation and cyclization of translucin III to form hyperforin in *H. perforatum*.**

**A**, LC–MS extracted ion chromatograms EIC at m/z 537.3938 of the hyperforin standard, an *H. perforatum* plant extract and the ethylacetate extracts from yeast combinatory co-expression of candidate prenyltransferases and upstream metabolic genes for biosynthesis of translucin III. Only the co-expression of *HpPT4* with *HpCCL2*, *HpPKS2*, *HpPT1*, *HpPT2* and *HpPT3* results in a distinct new peak, corresponding to the hyperforin standard.

**B**, Comparative MS/MS fragmentation analysis of the hyperforin standard, the corresponding compound in an *H. perforatum* extract and the product from the HpPT4-characterization experiment confirming the identity of the prenylated product as hyperforin.

**C**, Schematic of the pathway illustrating the prenylation and cyclization of translucin III to generate hyperforin catalyzed by HpPT4.

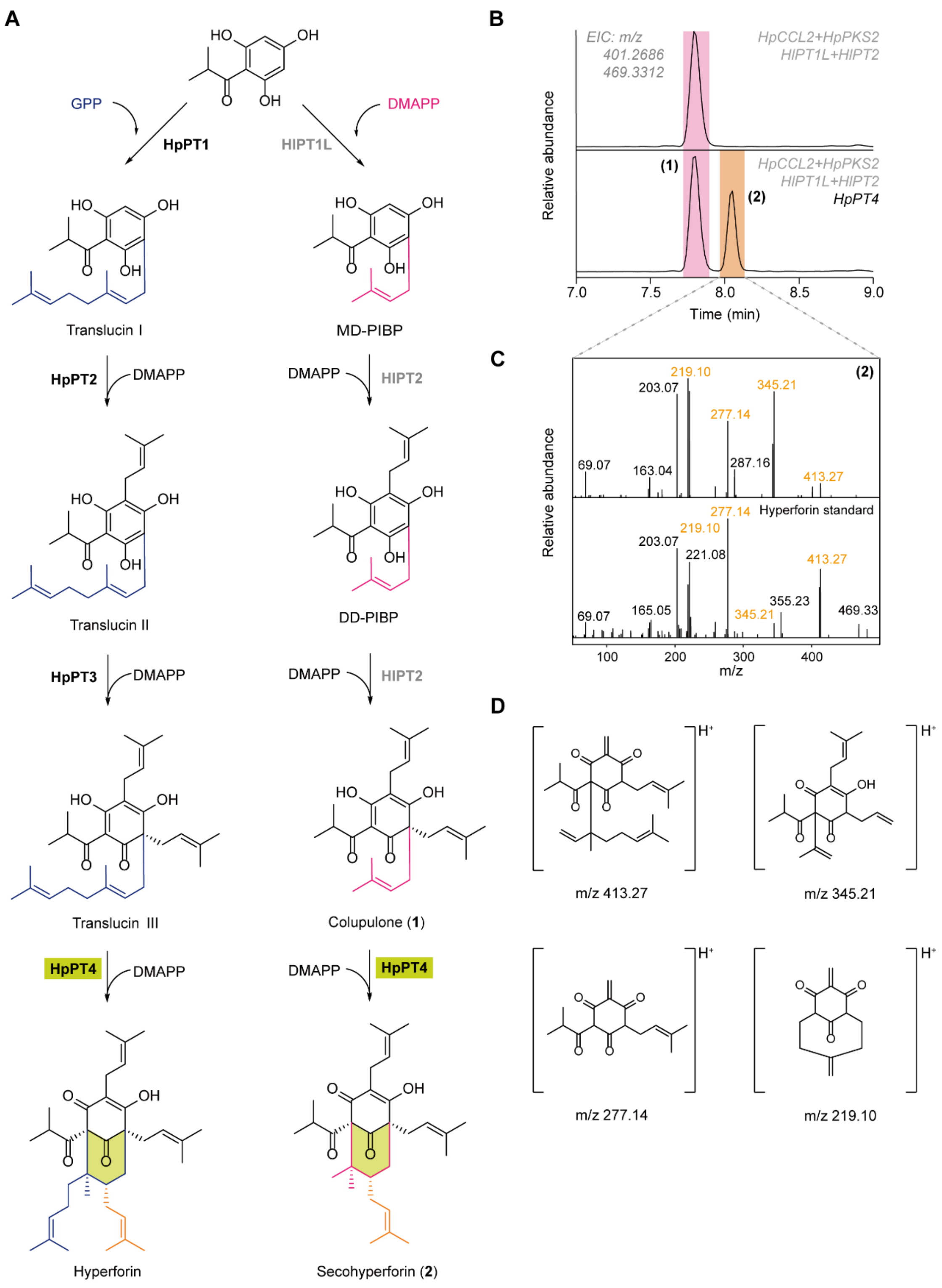


**Figure S16. Head-to-Middle Prenylation and Cyclization by HpPT4 in polycyclic polyprenylated acylphloroglucinol (PPAP) biosynthesis.**

**A**, The analogy between the biosynthesis of hyperforin and its reported analog secohyperforin (**2**) from *H. perforatum* cell cultures[12]. To test the potential role and substrate promiscuity of HpPT4 catalyzing the 1'-2 irregular prenylation and cyclization, we co-expressed in yeast the following combination: (a) *HpCCL2* and *HpPKS2* from *H. perforatum* for biosynthesis of PIBP[11]; (b) *HlPT1L* and *HlPT2* from hops (*Humulus lupulus*) for prenylation of PIBP and biosynthesis of colupulone (**1**)[13], an analog of translucin III. HpPT4 catalyzed the 1’-2 prenylation of colupulone (**1**) and cyclization to create secohyperforin (**2**) in similar fashion as it does in hyperforin biosynthesis.

**B**, EIC LC–MS chromatogram at m/z 401.2686 and 469.3312 highlights the emergence of a new peak secohyperforin (**2**), observable when *HpPT4* is co-expressed with *HlPT1L* and *HlPT2* from *H. lupulus*. Notably, HlPT1L and HlPT2 alone lead to the formation of colupulone (**1**).

**C**, Comparison of MS/MS spectra between (2) and hyperforin, demonstrating the similarities of $MS^2$ spectra indicating the similar $MS^2$ fragmentation reaction and structures.

**D**, Chemical structures of potential fragments identified in the MS/MS spectra corresponding to panel C[12].

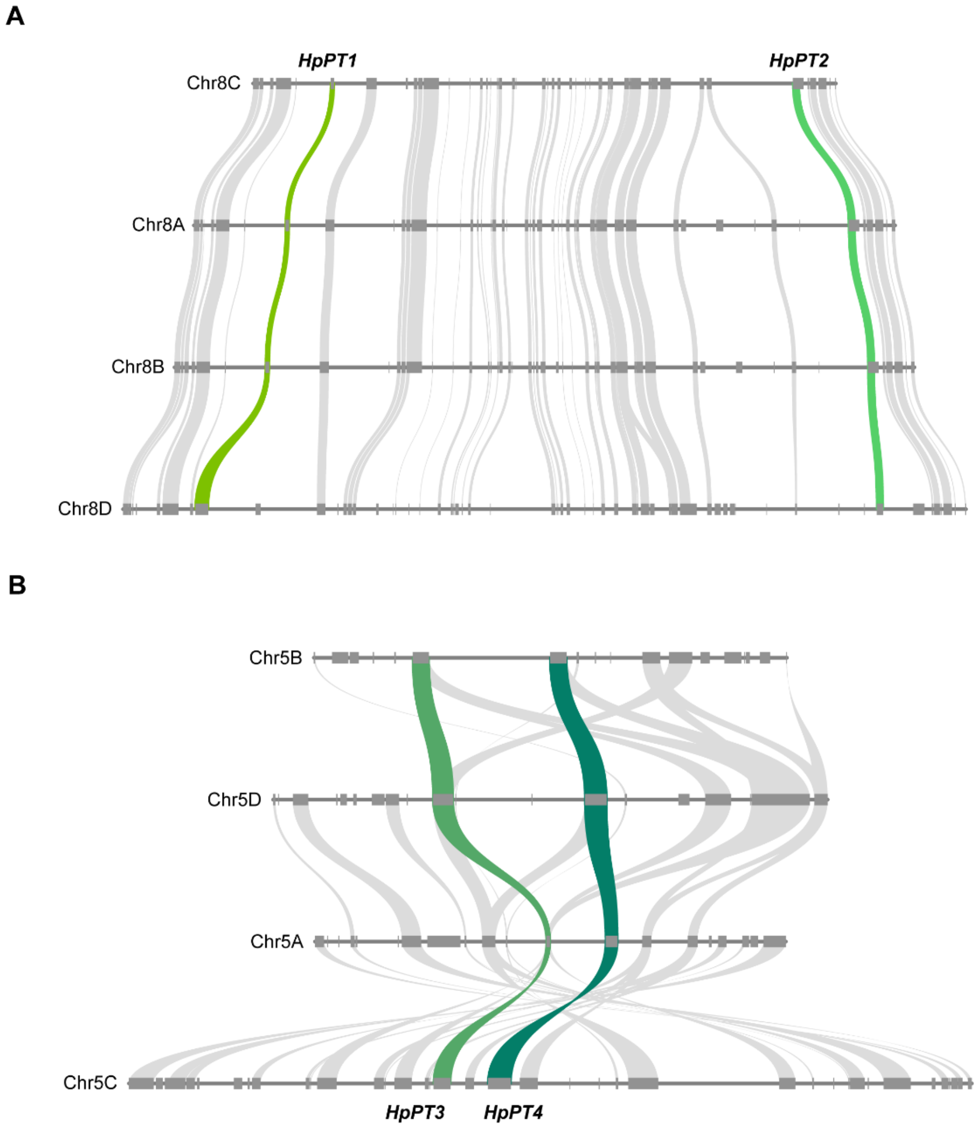


**Figure S17. Microsynteny analysis of prenyltransferases *HpPT1*, *HpPT2*, *HpPT3* and *HpPT4* across homologous chromosomes in the *H. perforatum* genome.**

**A**, Green ribbons illustrate the microsynteny of *HpPT1* and *HpPT2*, demonstrating the shared genomic locations on their respective homologous chromosomes.

**B**, Green ribbons illustrate the microsynteny of *HpPT3* and *HpPT4*, demonstrating the shared genomic locations on their respective homologous chromosomes.

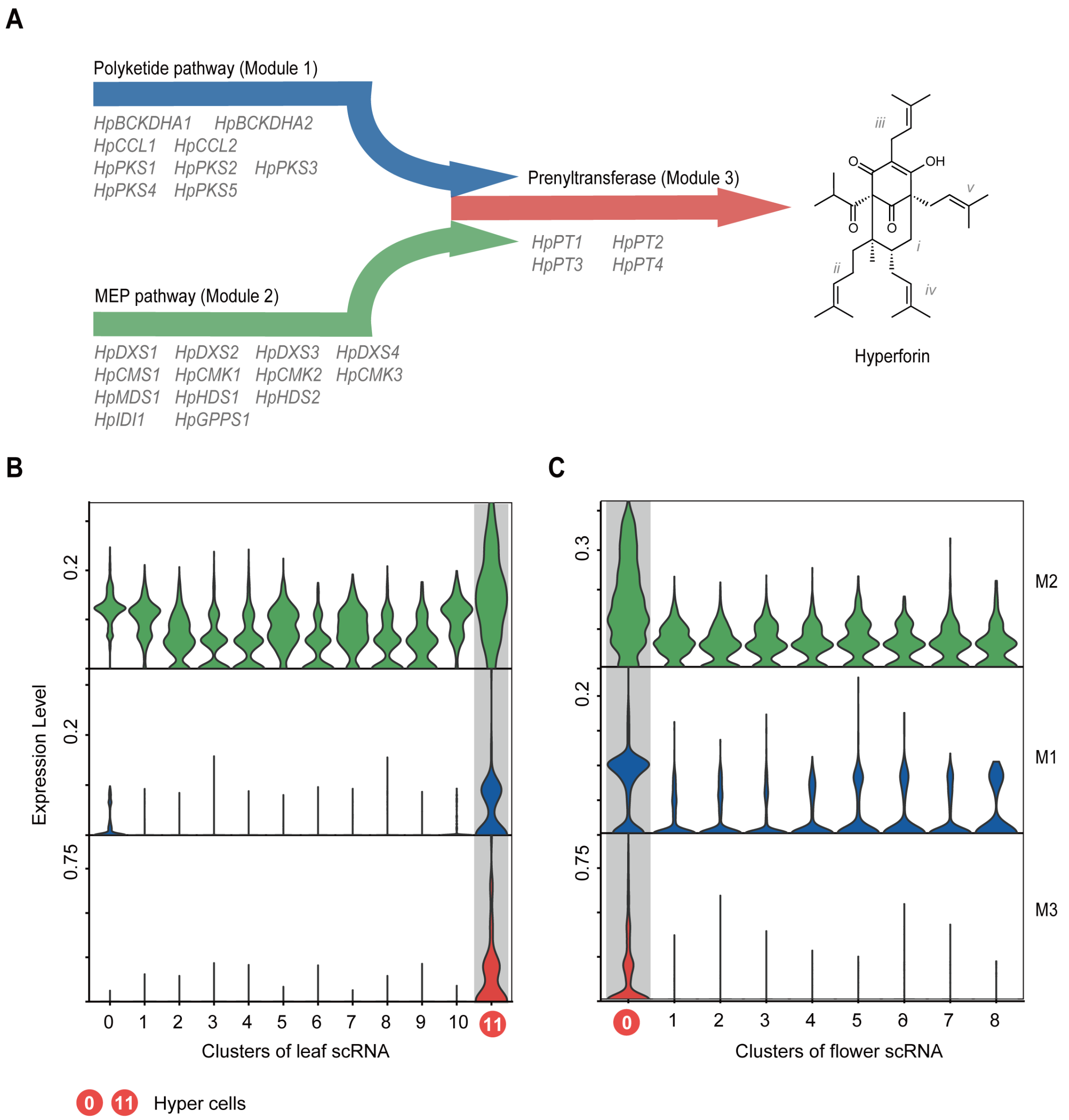


**Figure S18. Elevated gene expression in the three modules of hyperforin biosynthesis *de novo* in Hyper cells of *Hypericum perforatum*.**

**A**, Illustrative scheme of the three distinct modules (module 1: polyketide pathway – supplying the prenyl receptor PIBP; module 2: MEP pathway – supplying prenyl donors GPP and DMAPP; module 3: prenylation) involved in the biosynthetic pathway of hyperforin, including the associated metabolic genes, delineating the stepwise process leading to hyperforin formation.

**B-C**, Comparative expression profiles demonstrating heightened gene expression across all three biosynthetic modules in Hyper cells, as observed in both leaf (B) and flower (C) atlases.

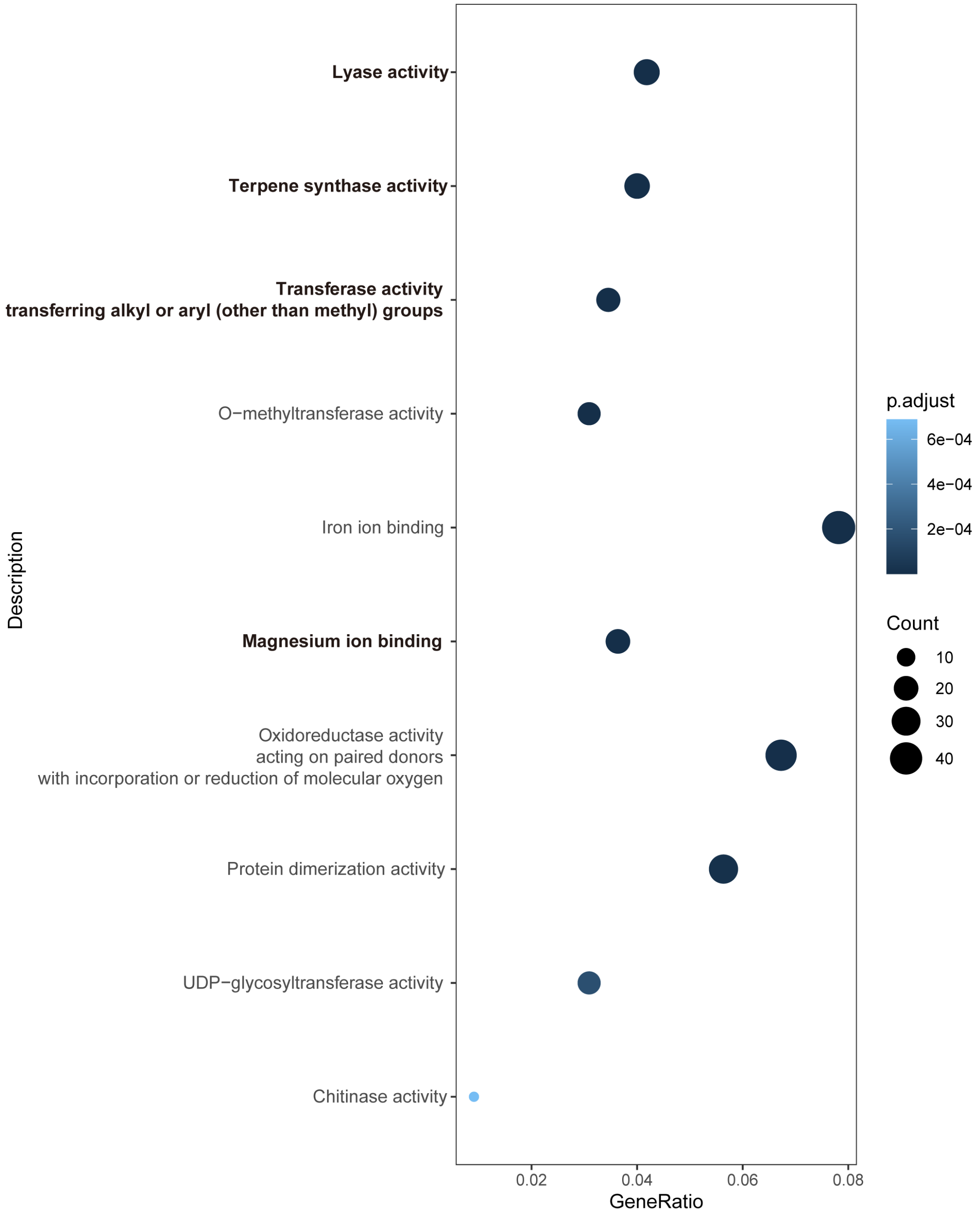


**Figure S19. GO enrichment analysis of up-regulated genes after methyl jasmonate treatment.** Two-months old seedlings of *H. perforatum* were sprayed with 400 μM MeJA. After 24 hours of treatment, the whole plants of seedlings with and without treatment were collected for further analysis.

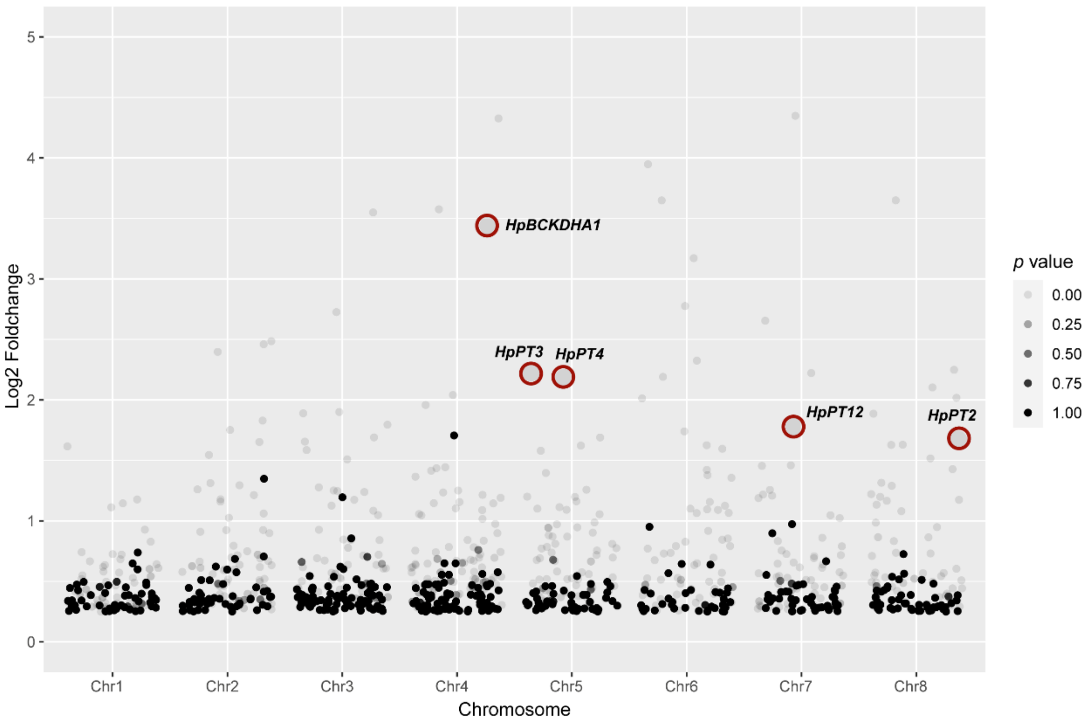


**Figure S20. Manhattan plot highlighting Hyper-cell marker genes in the *H. perforatum* leaf atlas (Table S12).**

Genes in the hyperforin biosynthesis pathway *HpBCKDHA1*, *HpPT2*, *HpPT3* and *HpPT4* are marked in red dots, indicating a high fold change in expression compared to other cell types.

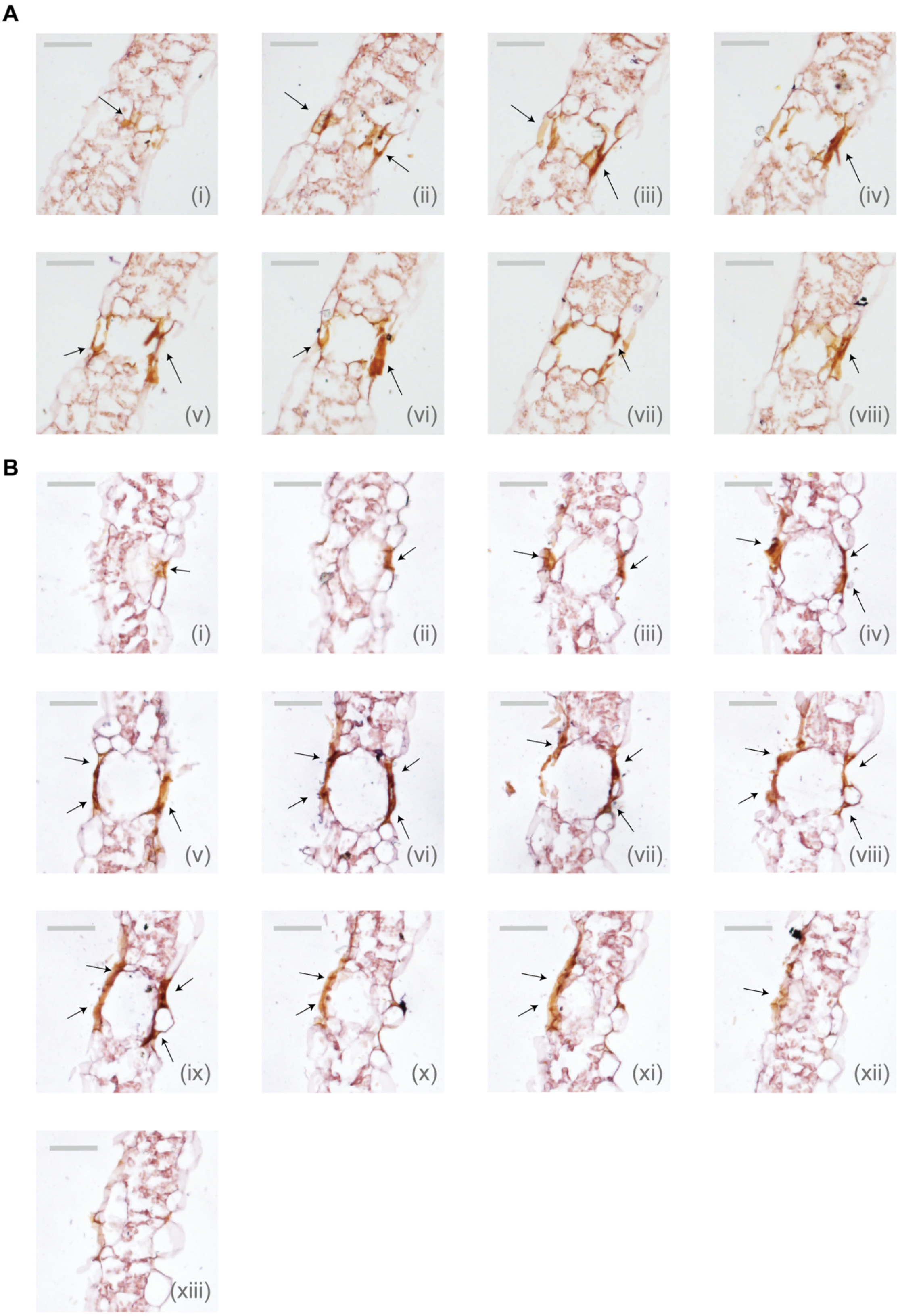


**Figure S21. RNA *in situ* hybridization of *HpPT3* expression in leaves.**
**A, B**, Consecutive cross-sections of *H. perforatum* leaves (two independent slices A and B), hybridized with probes derived from the conserved sequence of *HpPT3*. The arrows show the cellular location of the hybridization signals. Scale bars = 50 μm.

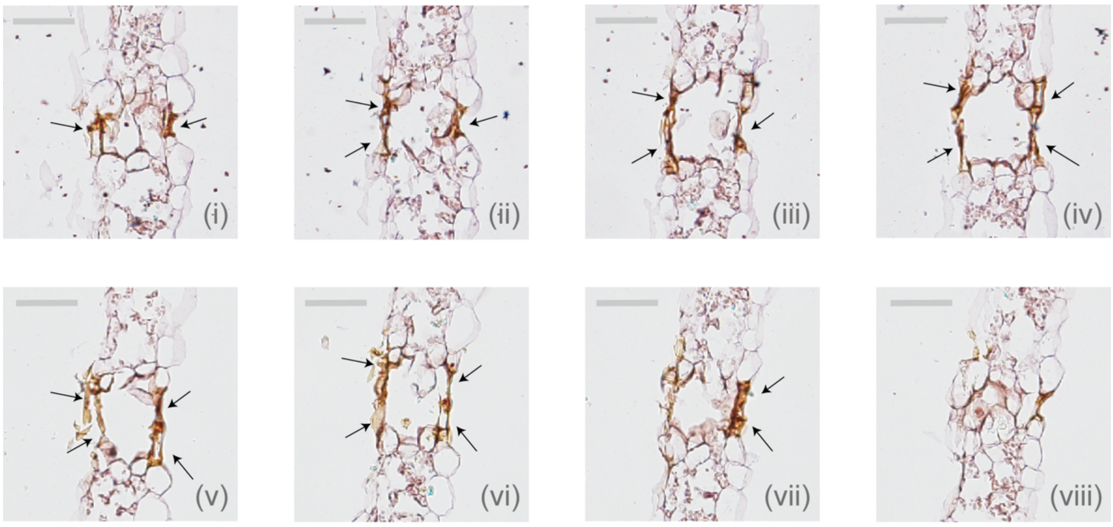


**Figure S22. RNA *in situ* hybridization of *HpPT4* expression in leaves.**

Consecutive cross-sections of *H. perforatum* leaves hybridized with probes derived from the conserved sequence of *HpPT4*. The arrows show cellular localization of the hybridization signals. Scale bars = 50 μm.

A

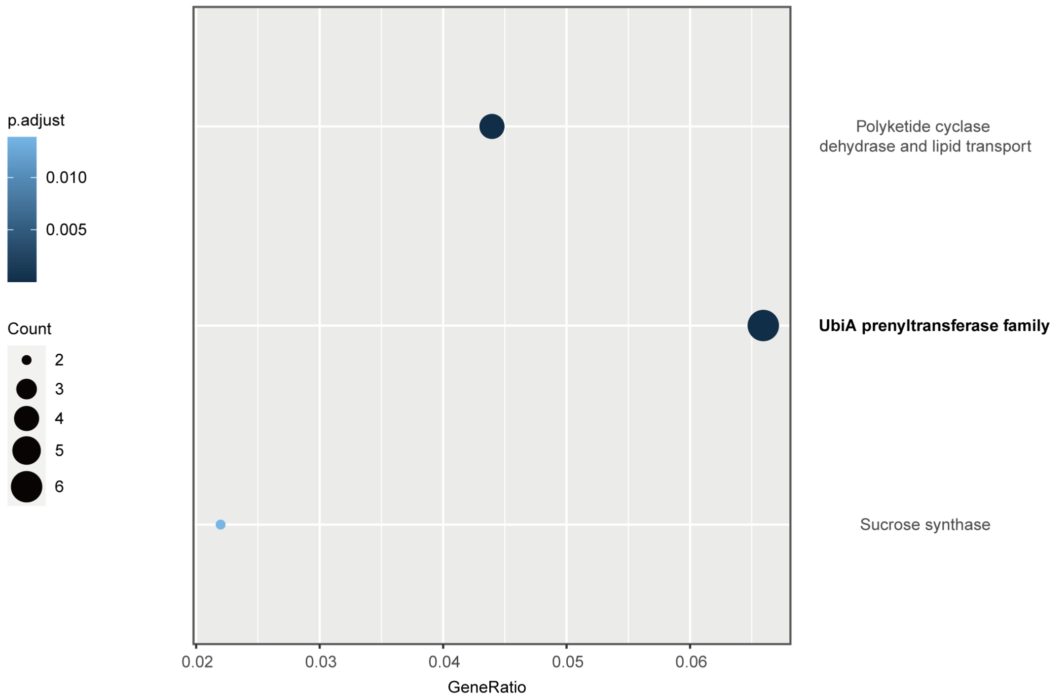


B

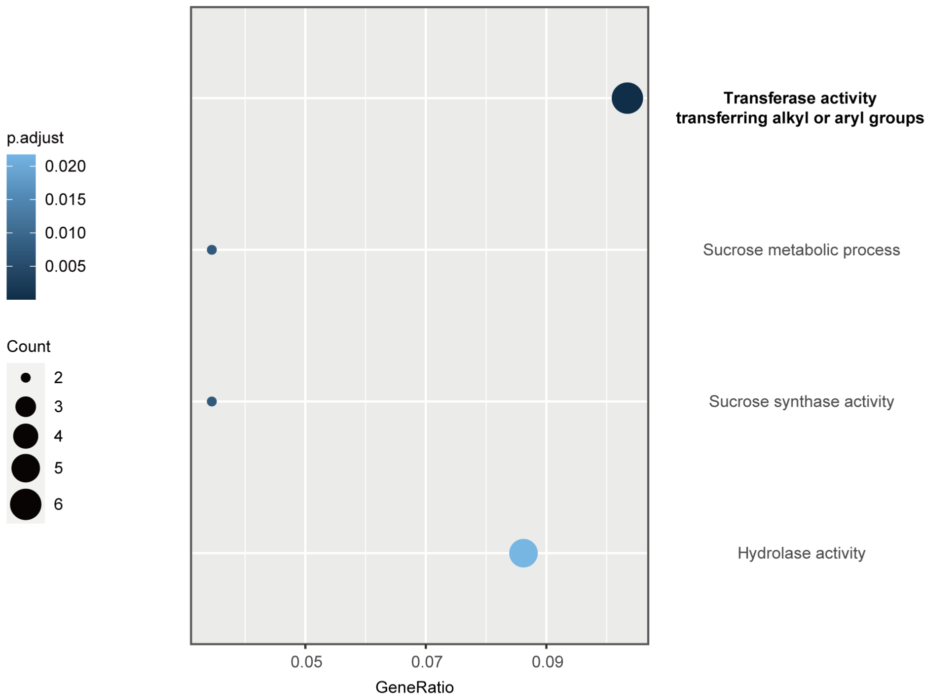


**Fig. S23. Enrichment Analysis of 'Hyper Cells' in the *H. perforatum* Flower Atlas.** **A**, Pfam domain enrichment analysis for 'Hyper Cells' indicates that the 'UbiA prenyltransferase family' ranks as the second most significantly enriched term. **B**, Gene Ontology (GO) term enrichment analysis highlights 'transferase activity' as the most prominent term.

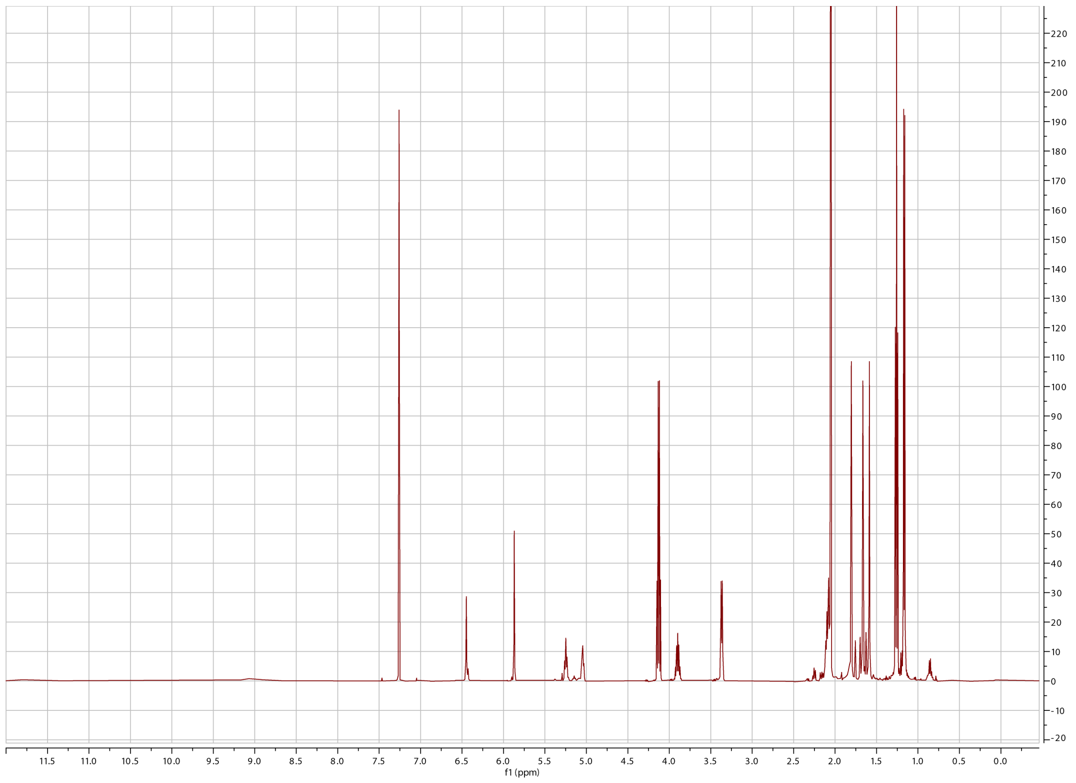


**Fig. S24. $^1$H-NMR (500 MHz) spectrum of translucin I in $CHCl_3$-$d_1$.**

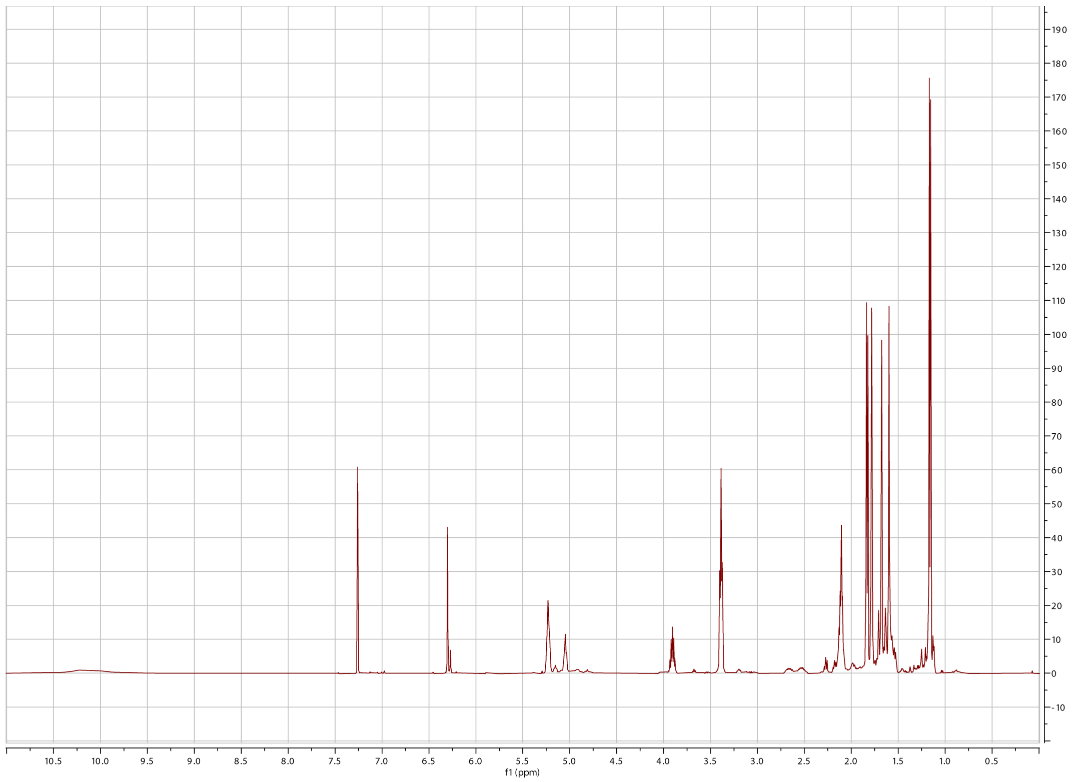

**Fig. S25.** $^{1}$**H-NMR (500 MHz) spectrum of translucin II in** $CHCl_3$-$d_1$.

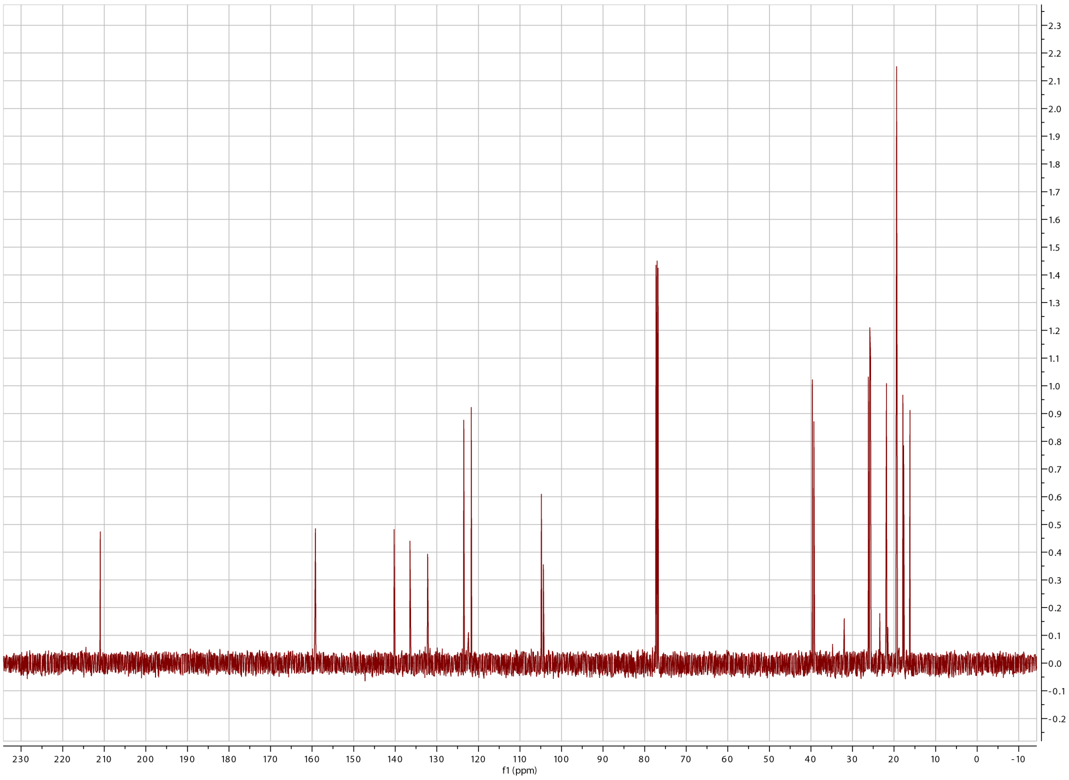


**Fig. S26.** $^{13}$**C-NMR (126 MHz) spectrum of translucin II in** $CHCl_3$-$d_1$.

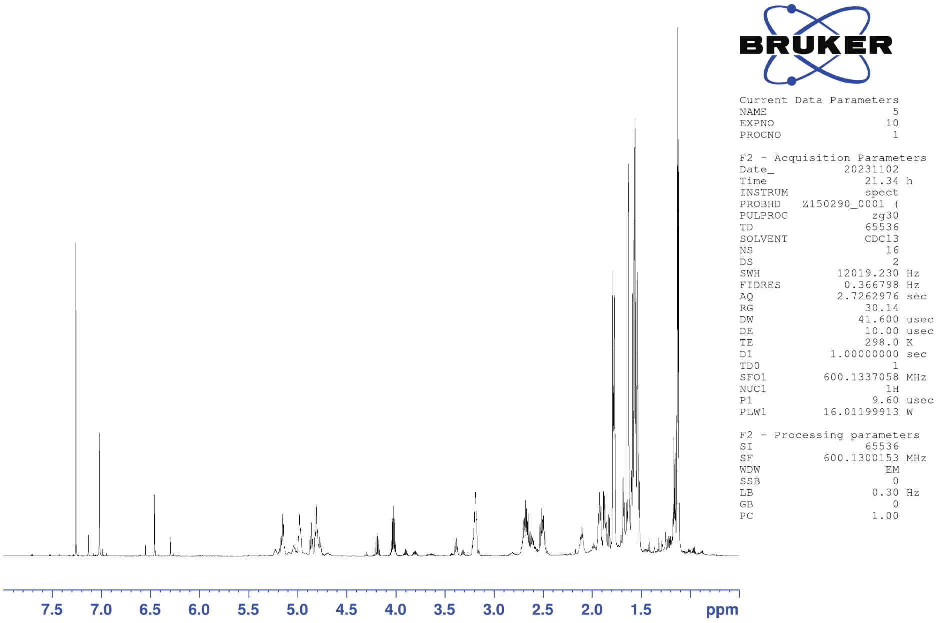


**Fig. S27. $^{1}$H-NMR (600 MHz) spectrum of translucin III in $CHCl_3$-$d_1$.**

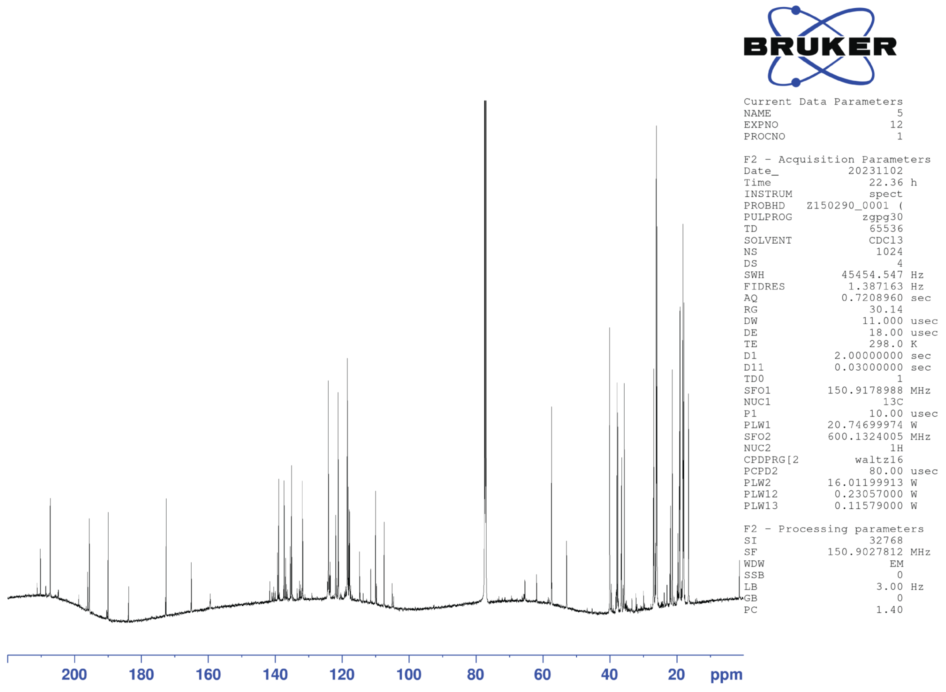


**Fig. S28.** $^{13}$**C-NMR (151 MHz) spectrum of translucin III in** $CHCl_3$-$d_1$.

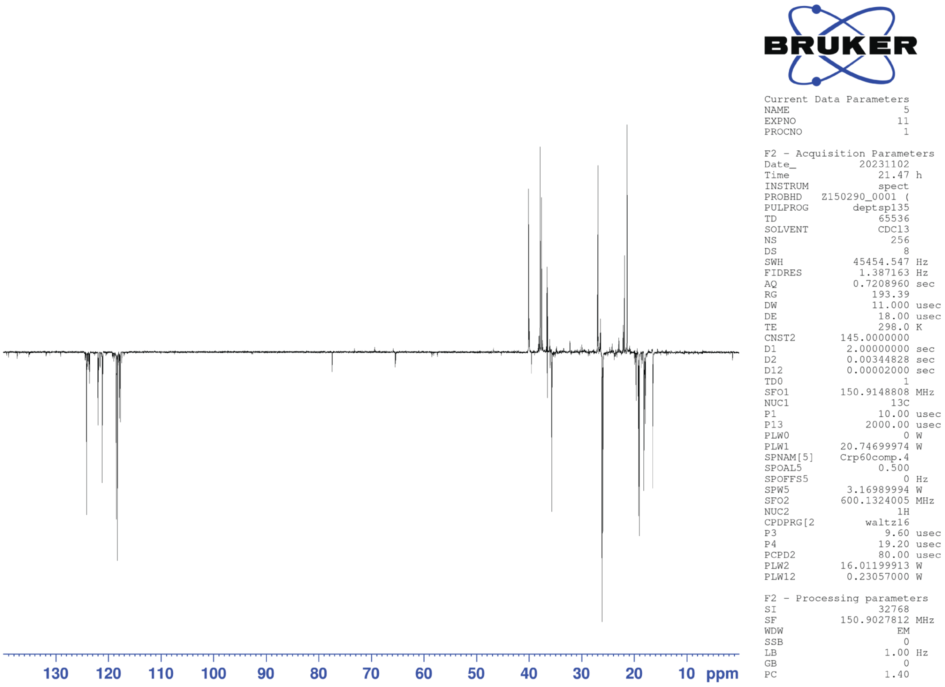


**Fig. S29. DEPT-135 $^{13}$C-NMR (151 MHz) spectrum of translucin III in $CHCl_3$-*d*$_1$.**

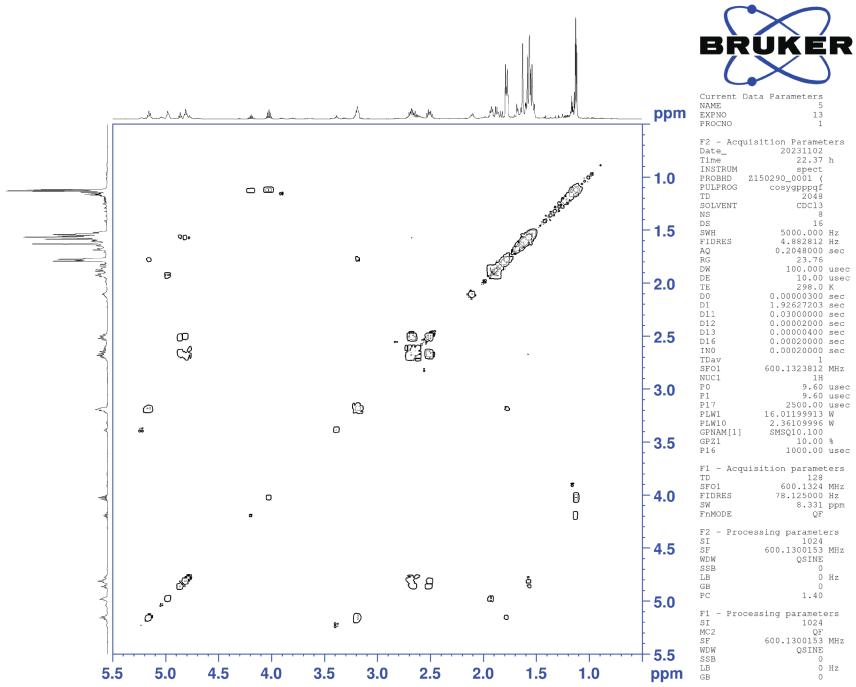


**Fig. S30. COSY spectrum of translucin III in $CHCl_3$-*d*$_1$.**

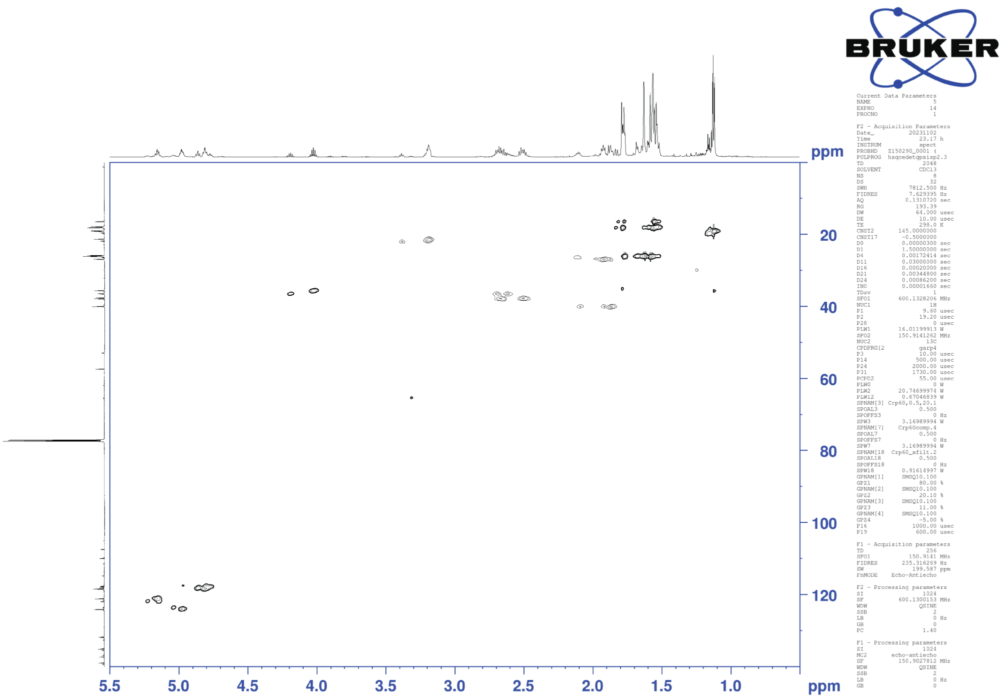


**Fig. S31. HSQC spectrum of translucin III in $CHCl_3$-$d_1$.**
